# Genetic drift does not sufficiently explain patterns of electric signal variation among populations of the mormyrid electric fish *Paramormyrops kingsleyae*

**DOI:** 10.1101/154047

**Authors:** Sophie Picq, Joshua Sperling, Catherine J. Cheng, Bruce A. Carlson, Jason R. Gallant

## Abstract

The mormyrid fish species *Paramormyrops kingsleyae* emits an electric organ discharge (EOD) with a dual role in communication and electrolocation. Populations of *P. kingsleyae* have either biphasic or triphasic EODs, a feature which characterizes interspecific signal diversity among the *Paramormyrops* genus. We quantified variation in EODs of 327 *P. kingsleyae* from 9 populations throughout Gabon and compared it to genetic variation estimated from 5 neutral microsatellite loci. We found no correlation between electric signal and genetic distances, suggesting that EOD divergence between populations of *P. kingsleyae* cannot be explained by drift alone. An alternative hypothesis is that EOD differences are a cue for assortative mating, which would require *P. kingsleyae* be capable of differentiating between divergent EOD waveforms. Using a habituation-dishabituation assay, we found that *P. kingsleyae* can discriminate between triphasic and biphasic EOD types. Nonetheless, patterns of genetic and electric organ morphology divergence provide evidence for hybridization between signal types. Although reproductive isolation with respect to signal type is not absolute, our results suggest that EOD variation in *P. kingsleaye* has the potential to serve as a cue for assortative mating and point to selective forces rather than drift as important drivers of signal evolution.

## Introduction

The evolution of diversity in courtship signals has important implications for mate recognition, mate preference and speciation, as evidenced by numerous rapidly evolved groups of species with highly diverse courtship signals (Diamond 1986; Allender et al. 2003; Mendelson and Shaw 2005b; Boul et al. 2006; Mullen et al. 2007; Arnegard et al. 2010b). Despite this well-established relationship, it is still a challenge in speciation research to understand the processes driving divergence in courtship signals and their consequences in the early evolution of reproductive isolation (Marie Curie SPECIATION Network 2012). Sexual selection has been proposed as the primary driver of signal divergence in a diversity of taxa, including frogs (Boul et al. 2006; Lemmon 2009), birds (Irwin et al. 2001; Toews and Irwin 2008) and crickets (Gray and Cade 2000; Mendelson and Shaw 2005a; Grace and Shaw 2011) (reviewed in Wilkins et al. (2013)). Detailed understanding of the mechanisms of signal divergence are well understood in only a few model systems (e.g. Maan et al. 2010; Chung et al. 2014; Ryan and Guerra 2014), partly owing to difficulties in understanding the anatomical and physiological substrates of signal generation and perception. In systems in which these substrates are at least partly understood, many signaling modalities are constrained by environmental effects on signal transmission and reception (Bradbury and Vehrencamp 2011), resource competition (Kingston and Rossiter 2004), or predation (Ryan 1988), making it difficult to disentangle the relative effects of sexual and natural selection. Signal divergence may also simply result from drift (Lynch and Hill 1986; Endler 1993), the fundamental alternative to selective forces, whereby stochastic variation accumulates in isolated populations over time (Irwin et al. 2008; Amézquita et al. 2009; Campbell et al. 2010; Picq et al. 2016).

Weakly electric fish such as African mormyrids are well suited for overcoming these experimental difficulties. Mormyrids communicate and electrolocate using electric signals termed electric organ discharges (EODs). These signals are diverse and are easily measured and quantified; importantly, they have a well-characterized and discrete anatomical and physiological basis related to the function of other electrically excitable tissues such as muscle and nerve (Bennett and Grundfest 1961; Arnegard et al. 2010a; Gallant et al. 2011; Carlson and Gallant 2013). Mormyrids represent a rapidly radiating family within the very ancient order of bony-tongued fishes (Osteoglossiformes) (Carlson and Arnegard 2011; Rabosky et al. 2013). There are at least two species radiations of African mormyrids, one in the genus *Campylomormyrus* (Feulner et al. 2007; Feulner et al. 2008) and a second in the genus *Paramormyrops* (Sullivan et al. 2002; Sullivan et al. 2004). A total of 11 *Paramormyrops* species are presently recognized, and more than 10 species with provisional code names remain to be described, all of which have rapidly diverged in the geographically restricted drainage basins of West-Central Africa. *Paramormyrops* exhibit highly divergent EODs, which vary primarily in duration (0.5-8 ms), the number of phases, and polarity (Fig. 1; Sullivan *et al*. 2000). Recent work has demonstrated that the evolution of electric signals far outpaces divergence in morphology, size and trophic ecology, implying over broad taxonomic scales that *Paramormyrops* diversification may largely have been driven by evolution of electric communication (Arnegard et al. 2010b). Multiple, non-exclusive hypotheses have been proposed for shaping signal diversity in electric fishes (Crampton 2019), including sexual selection (Arnegard et al. 2005; Feulner et al. 2009a), adaptation to environments with different electroreceptive predator pressures (Stoddard 1999), differential prey detection abilities (Feulner et al. 2009b), reproductive character displacement (Crampton et al. 2011), and drift (Picq et al. 2016).

**Figure 1.**
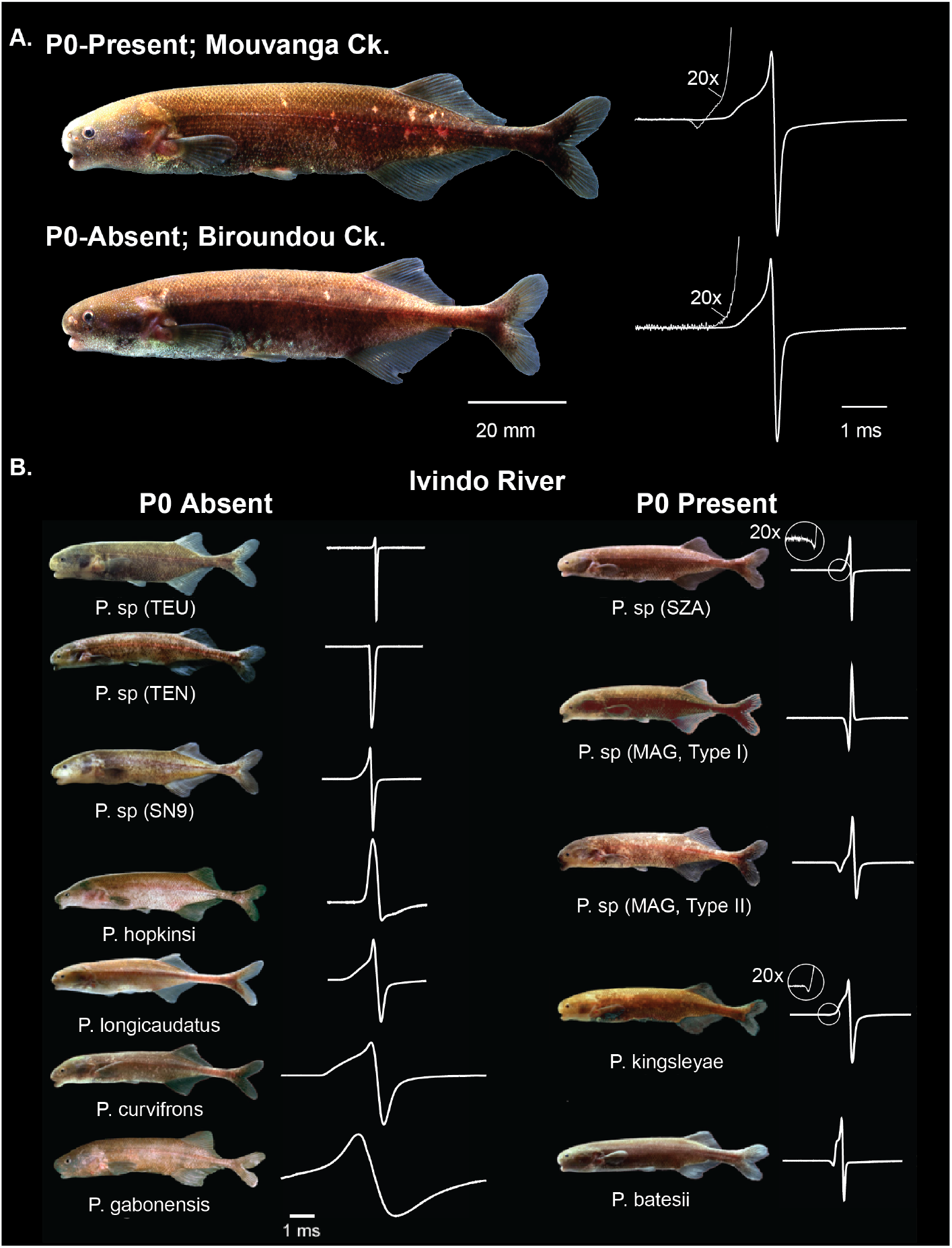
*Paramormyrops kingsleyae* EOD variation is a microcosm of EOD variation in *Paramormyrops*. **A**. *P. kingsleyae* EODs are variable in the presence of a small head negative phase (P0), see Gallant et al. (2011). **B**. 12 sympatric *Paramormyrops* specimens captured from the same locality in the Ivindo River illustrates EOD variation within *Paramormyrops* in terms of duration and polarity, as well as in the number of phases. All *Paramormyrops* on the left have P0-absent EOD waveforms, all *Paramormyrops* on the right have P0-present waveforms.

### The Study System

To better understand how broad macroevolutionary patterns in EOD trait divergence result from processes acting among individual populations, Gallant et al. (2011) examined patterns of geographic variation among the highly vagile, geographically widespread *Paramormyrops* species, *P. kingsleyae*. *P. kingsleyae* are polymorphic with regard to a small head-negative phase in the EOD, termed P0 (Fig. 1A). This signal feature is, at maximum, 2% of the peak-to-peak amplitude of some EODs, but is altogether absent in some geographically isolated forms. Despite this subtle difference in EOD signals, the anatomical substrate for P0-present and P0-absent signals requires a considerable structural reorganization of the cells that comprise the electric organ (electrocytes). Individuals with P0-present EODs possess electrocytes with penetrating stalks and anterior innervation (type *Pa*) whereas in P0-absent EODs, electrocytes have non-penetrating stalks with posterior innervation (type *NPp*). In this regard, *P. kingsleyae* represents a “microcosm” of signal diversity within the *Paramormyrops* genus, as this is also the main component of EOD divergence characterizing inter-species signal differences (Gallant et al. 2011). P0-present EODs from mormyrids with penetrating-stalked electrocytes originated early in the sub-family Mormyrinae. They are widespread among the 20 known genera, with at least eight separate presumed reversions to non-penetrating stalked electrocytes in various genera, with five additional independent reversions in *Paramormyrops* alone. (Fig. 5 in Carlson & Gallant 2013). The majority of *P. kingsleyae* populations across Gabon have P0-present EODs (Fig. 2; Gallant et al. 2011). However, in several populations within a watershed region of the Louétsi river in Southern Gabon, every individual has P0-absent EODs.

**Figure 2.**
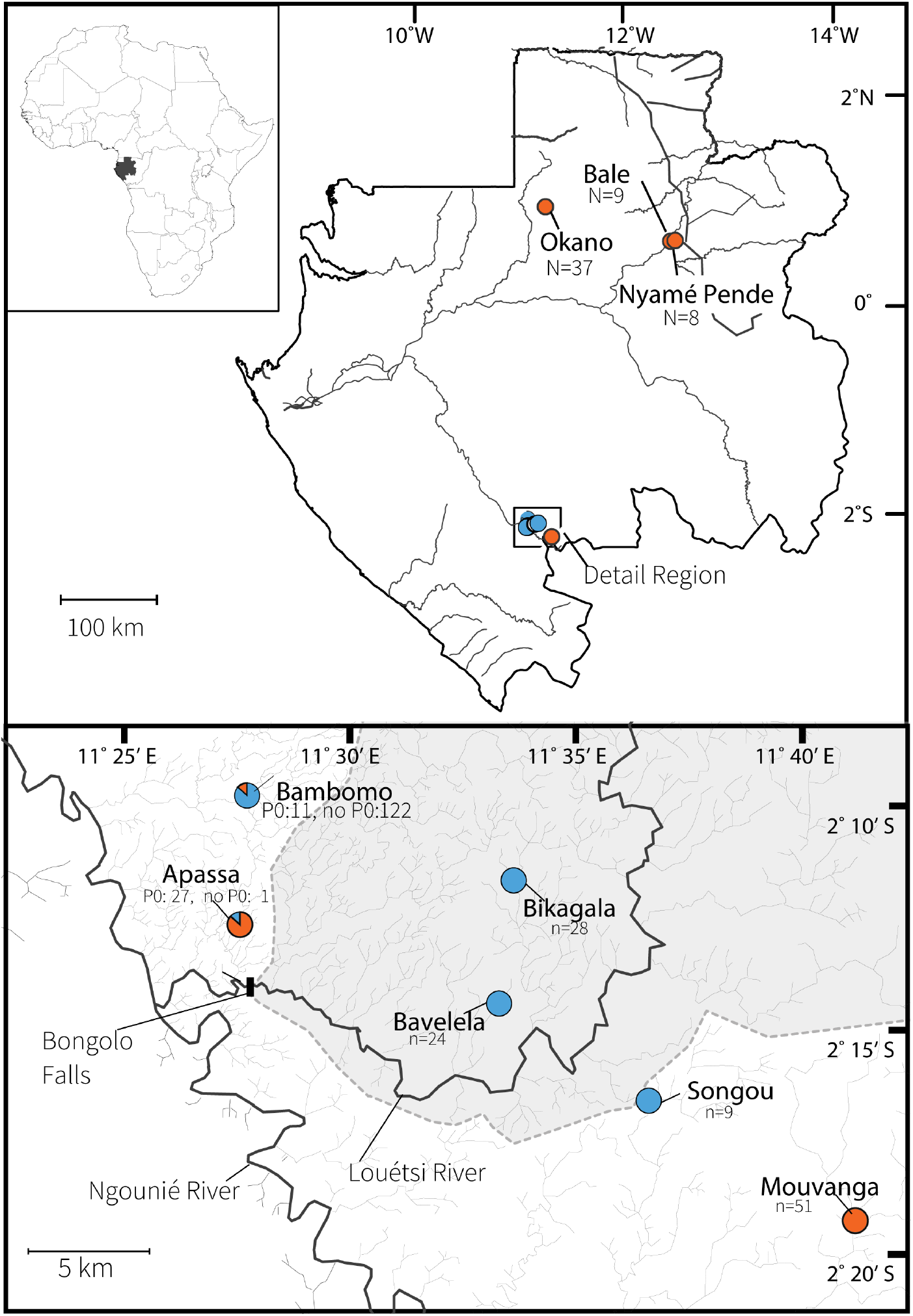
Map of study populations. Top map shows relationship between populations across Gabon. Box in the south indicates the location of the higher detail map, which indicates the relationship of Southern populations near the confluence of the Louétsi and Ngounié rivers. For all maps, populations are indicated by pie charts, as reported in Gallant et al. 2011. Orange indicates proportion of individuals P0-present, blue indicates proportion of individuals that were P0-absent. Bongolo falls is indicated in the map of southern populations, and a grey region bounded by a dotted line indicates the small streams and creeks that drain into the Louétsi river above Bongolo Falls.

Gallant et al. (2011) considered several alternative hypotheses to explain the observed geographic patterns of variation in *P. kingsleyae* EODs and considered drift as the most likely, based on three major findings: (1) *P. kingsleyae* exhibits clinal variation in EOD signals in duration and P0 magnitude; (2) the discovery of morphologically ‘intermediate’ individuals in Apassa and Bambomo Creek, where both biphasic and triphasic signal types co-occur, which were putatively considered as evidence of hybridization. This suggested that the ability to discriminate EOD waveforms within *P. kingsleyae* may not exist, or that mating may not be fully assortative with respect to signal type; (3) evidence from Arnegard et al. (2005) showing substantial genetic partitioning between allopatric populations of *P. kingsleyae* in southern Gabon, indicating limited gene flow between allopatric populations in the Louétsi drainage region. Taken together, these results suggest that signal divergence may increase with geographic distance, that mating may be random with respect to signal type in sympatric populations, and that allopatric *P. kingsleyae* populations are genetically isolated, patterns that are expected when drift is important in driving signal divergence (Wilkins et al. 2013).

In this study, we chose to explicitly test the hypothesis that genetic drift is responsible for the biogeographic distribution of EOD signals in *Paramormyrops kingsleaye*. First, we compared variation in EOD waveforms between *P. kingsleyae* populations to variation in neutral genetic markers in the same individuals. If the present biogeographic distribution of signal types is mainly the result of neutral processes, we would expect signal divergence to increase linearly with genetic divergence. Alternatively, if adaptive forces were the main determinants of signal diversity (Wilkins et al. 2013), we would expect no correlation between signal divergence and genetic divergence. Our results show that there is no relationship between signal divergence and genetic divergence, rejecting our hypothesis of neutral processes explaining phenotypic divergence among *P. kingsleyae*.

A prominent alternative hypothesis for the distribution of signals is that mormyrid species may mate assortatively on the basis of EOD differences (Arnegard et al. 2005; Feulner et al. 2009a, b; Arnegard et al. 2010b). For this to be a valid explanation for the present distribution of signal types within *P. kingsleyae*, it would require that *P. kingsleyae* have perceptual ability to discriminate EOD differences. We assessed this directly, using a dishabituation behavioral assay to determine whether *P. kingsleyae* from a sympatric location are able to discriminate between EOD types. We also assessed this hypothesis indirectly by examining patterns of genotypic differentiation and electric organ morphology to determine whether there was any evidence of assortative mating among *P. kingsleyae* EOD types.

## Materials and Methods

### Field collections

Collections of *Paramormyrops kingsleyae* specimens were made during field trips to Gabon, West Central Africa in 1999, 2001 and 2009 from 9 locations summarized in Table 1 and Figure 2. A catalog of individual specimens used in this analysis is provided as Table S1, which provides Cornell Museum of Vertebrates accession numbers for all specimens used in this paper, along with associated metadata. We sampled two locations, Mouvanga and Bambomo Creeks, twice: once in 1999-2001 and again in 2009. Samples collected in different years at these locations were kept separate for all analyses, resulting in a total of 11 sampled populations.

**Table 1.** Summary of locations and years of population samples, and analyses performed on each population. * Indicates additional individuals were sampled from these populations to perform these analyses. Numbers in parentheses indicate sample sizes, if no sample size is reported, the entire N reported was analyzed. Column EO indicates population utilized in histological analysis of electric organs (see methods). Column Behavior indicates population utilized in EOD discrimination task (see methods). Table S1 contains the full metadata for all specimens, including Cornell Museum of Vertebrates accession numbers for vouchers is provided.

| **Pop** | **Year** | **Lat** | **Long** | **N** | **Genotype** | **EOD** | **EO** | **Behavior** |
| --- | --- | --- | --- | --- | --- | --- | --- | --- |
| Apassa | 2009 | 2° 12' 42" S | 11° 27' 50" E | 28 | **X** | **X** | **X (4)** |  |
| Bale | 2001 | 0° 31' 9" S | 12° 47' 58" E | 8 | **X** | **X** |  |  |
| Bambomo | 2009 | 2° 9' 49" S | 11° 27' 42" E | 106 | **X** | **X** | **X (7)** | **X (17)\*** |
| Bambomo | 1999 | 2° 9' 49" S | 11° 27' 42" E | 27 | **X** | **X** |  |  |
| Bavavela | 2009 | 2° 14' 33" S | 11° 33' 22" E | 24 | **X** | **X** |  |  |
| Bikagala | 2009 | 2° 11' 43" S | 11° 33' 40" E | 28 | **X** | **X** |  |  |
| Mouvanga | 2009 | 2° 19' 23" S | 11° 41' 18" E | 32 | **X** | **X** | **X (6)** | **X (10)\*** |
| Mouvanga | 1999 | 2° 19' 23" S | 11° 41' 18" E | 19 | **X** | **X** |  |  |
| Nyame Pende | 2001 | 0° 30' 16" S | 12° 47' 48" E | 9 | **X** | **X** |  |  |
| Okano | 2001 | 0° 48' 57" S | 11° 39' 1" E | 37 | **X** | **X** |  |  |
| Songou | 2009 | 2° 16' 42" S | 11° 36' 41" E | 9 | **X** | **X** | **X (4)** |  |

Fish were collected using a variety of methods, including fish traps baited with worms, hand nets combined with electric fish detectors, and by light rotenone applications. Following any application of rotenone, we immediately transferred the fish to fresh, aerated water, where they recovered completely.

We georeferenced sampling locations and calculated pairwise geographic distances between all study populations using digitized topographic maps, which were superimposed over satellite imagery provided in Google Earth software (Google, Inc. v.6.0.1). For each pair of populations, the distance between any two populations was assumed to be the shortest river path between the populations and was calculated by tracing currently mapped rivers between these populations. We note that additional, shorter paths connecting populations could potentially be created by seasonal flooding events (see Arnegard et al. 2006).

### EOD recordings

EODs of each specimen were originally recorded within 24 hours of capture in 1- to 5 liter plastic boxes filled with water from the collection site. Signals were recorded with bipolar chloride-coated silver wire electrodes and amplified (bandwidth =0.0001-50 kHz) with a differential bioamplifier (CWE, Inc: Ardmore, PA), and digitized at a 100 kHz-1 MHz sampling rate, with head-positivity plotted upward using a Daqbook or WaveBook (IOTECH: Cleveland, OH), or a USB-powered A-D Converter (National Instruments: Austin, TX). All EOD recordings were made at a vertical resolution of 16 bits per sample. After recording their EODs, we euthanized individual specimens with an overdose of MS-222. We removed one or more paired fins from specimens and preserved these tissues in 95% ethanol. Each specimen was given a unique specimen identification tag, and fixed free-floating in 10% formalin (phosphate-buffered; pH 7.2) for at least 2 weeks. Specimens were then transferred to 70% ethanol and deposited in the Cornell University Museum of Vertebrates. All methods conform to protocols approved by Cornell University’s Center for Research Animal Resources and Education.

### Analysis of electric signal variation

Following the methods described by Gallant *et al*. (2011) and by Arnegard *et al*. (2003), we made 21 measurements (Fig. 3, Table S2) from each recorded EOD waveform using a custom written program in MATLAB (Mathworks, Inc.: Natick, MA). For each waveform, we calculated amplitudes, times, and slopes at nine landmarks defined by peaks, zero crossings, first derivative peaks, and threshold crossings (Fig. 3; Table S2). In addition, we calculated a power spectrum for each EOD waveform using the MATLAB *Fast Fourier transform* function to determine the peak frequency and a low and high frequency with power 3 dB below the peak frequency for each EOD recording. We quantified patterns of EOD variation among all the *P. kingsleyae* individuals collected by performing principal components analysis (PCA) on the set of all 21 measurements (normalized by using the function *scale in* R version 3.4.3 (R Core Team 2017) using the R function *princomp*. Electric signal distance between populations was calculated as the Euclidean distance between group centroids for the first two principal components using the R package *vegan* (Oksanen et al. 2018). To test the null hypothesis of no differentiation between EOD waveforms from different populations, we also computed the Euclidean distance between all pairs of individuals and performed a permutational MANOVA (PERMANOVA) directly on this pairwise signal distance matrix using 1000 permutations to determine probability values, using the *adonis* function from *vegan*.

**Figure 3.**
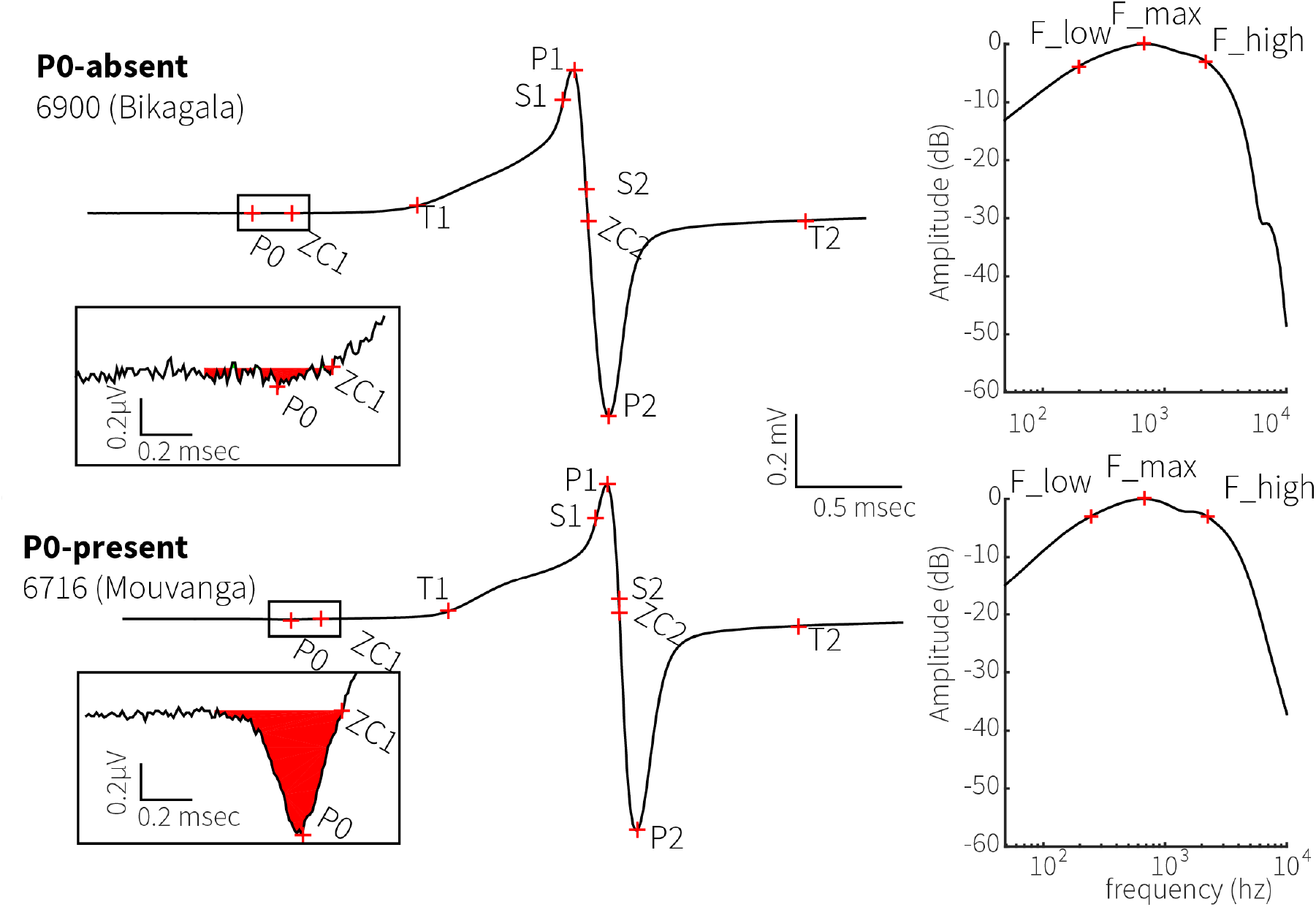
Example EOD Waveforms and Landmarks Measured. EOD landmarks and measurements used in the PCA were identical to those described in Gallant *et al*. (2011) and are listed for the reader’s convenience as Table S2. We indicate two example EOD waveforms: P0-absent (**A**) and P0-present (**B**) to illustrate the locations of these landmarks on EOD waveforms and their associated power spectra. Voltages shown are relative to a normalized P1-P2 voltage set equal to 1.0 volts, as described in methods.

Landmark-based signal processing methods as described above focus only on pre-selected signal features that do not encompass a comparison of the entire waveforms and may thus be discriminatory. Therefore, we complemented our PCA analysis by performing cross-correlation analysis of all 327 waveforms, as described in Carlson et al. (2011) and performed in Picq et al. (2016). EOD waveforms that were sampled at 150, 200, or 250kHz (n = 57) were first down-sampled to 100kHz using the *downsample* and *resample* functions in MATLAB. We then used the maximum of the absolute values of the cross-correlation function as a measure of pairwise waveform similarity, resulting in a matrix of pairwise similarities ranging from 0 (dissimilar waveforms) to 1 (identical waveforms). Multidimensional scaling (MDS) was then applied to this cross-correlation matrix using the “mdscale” function in Matlab with Kruskal’s normalized stress 1 criterion (Kruskal and Wish 1978). Electric signal distance between populations was calculated as the Euclidean distance between group centroids in the MDS space using *vegan.* Electric signal distances between all pairs of individuals were also estimated in the same way. As for the PCA, a PERMANOVA analysis was also performed on the individual pairwise signal matrix to test the null hypothesis of no differentiation between EOD waveforms from different localities.

Recording temperature is known to affect EOD duration (Kramer and Westby 1985). Gallant et al. (2011) showed that applying a Q10 temperature correction on 491 *P. kingsleyae* EODs recorded across Gabon at temperatures ranging from 21 to 26.7°C did not result in a significantly different PCA scores of signal variation. As the range of recording temperatures in the current study is narrower (20.5 to 25.1°C), we considered the effect of temperature on our PCA configuration of signal variation as negligible. Nevertheless, we still investigated the effect of temperature on signal variation quantified through multidimensional scaling of signal cross correlations. The full methodology and results of this analysis are included in Appendix 1. We concluded from this analysis that recording temperature was not a significant source of variation in our study, and therefore present our analysis on the full dataset without temperature correction for both PCA-derived and MDS-derived signal analyses.

### Microsatellite genotyping

We extracted DNA from the ethanol-preserved fin clips using DNeasy Tissue Kits (Quiagen, Inc.) for the 1998-1999 samples, or AgenCourt DNAadvance kits (Beckman-Coulter, Inc) for the 2009 samples. The 1998-1999 DNA samples originally genotyped by Arnegard et al. (2005) were re-genotyped for this study so that genotypes could be scored using identical criteria. We amplified DNA at each of five microsatellite loci (NBB001-NBB005) originally identified by Arnegard et al. (2005) using the Quiagen Type-It multiplex PCR System (Quiagen, Inc.). Reaction volumes were 15µl, consisting of 1µl template DNA, 7.5µl Type-it Multiplex Master Mix (containing HotStarTaq Plus DNA Polymerase and PCR buffer with 6µM MgCl_2_), and 2 pmol of each primer (5’ primers labeled with Applied Biosystems fluorescent dyes FAM, HEX or NED). Thermal cycling (under mineral oil) was 5 min at 95°C (initial activation) followed by 28 cycles of 95° for 30 sec, 60°C for 90 sec, and 72°C for 30 sec. Each individual reaction (containing PCR products for all 5 loci) was resolved by electrophoresis on an ABI 3100 automated sequencer (Applied Biosystems). Under these thermal conditions, reactions for locus NBB004 failed for the Bambomo, Nyamé-Pendé, and Balé Creek populations. For these populations, an additional PCR reaction was successfully performed as above using primers only for the NBB004 locus, with thermal cycling (under mineral oil) for 5 min at 95°C (initial activation) followed by 35 cycles of 95° for 30 sec, 50°C for 90 sec and 72°C for 30 sec. Following genotyping, individual fragment lengths were analyzed and binned according to size by visual inspection, using Genemapper 4.1 software (Applied Biosystems).

### Genotyping data analysis

For each microsatellite locus (NBB001-NBB005), we examined possible deviations from expected Hardy-Weinberg equilibrium within populations using the two-tailed exact test (Weir 1990) as implemented by GENEPOP v4.1 (Rousset 2008). Next, we performed exact tests of linkage disequilibrium between all pairs of loci (within and between populations), to test the independent assortment of loci. Statistical significance in both sets of tests was evaluated using Markov chain methods (10,000 dememorization steps; 1,000 batches; 5,000 iterations per batch). We additionally calculated observed and expected heterozygosity under Hardy-Weinberg equilibrium for each locus in every population using the software Arlequin 3.5 (Excoffier and Lischer 2010). The significance of each was evaluated at both the p=0.05 α probability threshold and the Bonferroni corrected threshold of p=0.001.

### Population structure

We quantified genetic differentiation between populations using F_st_ (Weir, Cockerham, 1984) with Arlequin v 3.5 (Excoffier and Lischer 2010). This standardized measure of population genetic structure is equivalent to the variance of allele frequencies between populations (i.e., subpopulations, or s) divided by the variance of allele frequencies in the total population consisting of both subpopulations combined (t). Using the same software, we evaluated the statistical significance of the F_st_ estimates (at both the uncorrected and Bonferroni-corrected thresholds) by permuting genotypes between populations 50,000 times.

Because only five microsatellite loci were scored, we performed power analysis simulations using POWSIM version 4.1 (Ryman and Palm 2006) to determine if our sample sizes, number of microsatellite loci, and allele diversity were sufficient to detect genetic differentiation (see full methodology and results in Appendix 2). These analyses showed that our sample sizes and specific genetic markers were adequate for detecting levels of genetic differentiation as low as F_st_ = 0.003 with a high probability (>0.9) (Fig. A2). We can therefore consider our genetic markers as sufficiently powerful for the purpose of this study, namely to give a reliable estimate of genetic differentiation between *P. kingsleyae* allopatric populations.

### Isolation by distance

Genetic isolation by distance (IBD) was explored following Rousset (1997), *i.e.* using a linear regression between genetic distance (F_st_/(1-F_st_)) versus geographic distance for all pairs of populations. The correlation was tested with a Mantel test (Mantel 1967), which accounts for non-independence of points in a distance matrix, using 1000 permutations in the R package *vegan*. The IBD analyses were performed on the whole dataset and repeated on Southern populations separated by short distances (<120 km) where sampling was high (including Bambomo, Apassa, Bikagala, Bavavela, Songou, and Mouvanga, Fig.2 lower panel). When considering the entire dataset, it is important to note that populations included in this study represent two distant and separate zones along the species’ otherwise continuous distribution across Gabon (Gallant et al. 2011). To control for this gap distribution in our sampling efforts and for the associated potential confounding effect of hierarchical structure (i.e. of regional-level effects between discontinuous Southern and Northern populations) on relationships between distance matrices, we ran a partial Mantel test, including a model matrix of regional membership that identified which populations were included in each pairwise comparison (e.g. North-North, South-South, South-North; (Smouse et al. 1986; Meirmans 2012). When only considering the subset of Southern populations, we additionally tested for the effect of Bongolo Falls as a barrier to gene flow between upstream (Bavavela and Bikagala) and downstream (Bambomo, Apassa, Songou, Mouvanga) populations by running a Mantel test between genetic distance and a model matrix coding for populations separated (1) or not separated (0) by this putative barrier.

### Comparison of signal and genetic divergence

To investigate whether electric signals evolve at a rate consistent with neutral evolution, we tested the correlation between signal and genetic distance for all pairs of populations using Mantel tests. This was performed for the whole dataset with a partial Mantel test (accounting for potential regional-level effects between discontinuous Northern and Southern populations) and repeated on Southern populations. We also tested whether signal differences between populations could be the product of isolation by distance by testing the correlation between signal and geographic distances, again using a partial Mantel test for the whole dataset and a Mantel test for the Southern populations. The potential effect of Bongolo Falls in driving signal divergence in Southern populations was tested using the same rationale as for the genetic data: a Mantel test was performed between signal distance and a model matrix coding for populations separated or not by the falls. All analyses were performed on both PCA and MDS derived signal distances.

### Evaluating discrimination ability: behavioral playback experiments

We performed two sets of electrical playback experiments to assess the ability of *P. kingsleyae* to discriminate P0-present and P0-absent waveforms. For all experiments, we assessed behavioral discrimination of EOD waveforms using a dishabituation paradigm described in detail by Carlson et al. (2011). Specimens were placed in a rectangular PVC enclosure containing both chlorided silver wire stimulus electrodes (Ag/AgCl) and Ag/AgCl recording electrodes, with the two pairs of electrodes oriented orthogonally with respect to one another. We delivered stimulus trains consisting of 10 bursts of 10 EOD pulses each, with an intra-burst interval of 30 ms, and inter-burst interval of 10 s, with a peak-to-peak intensity of 145 mV/cm. We constructed negative control trains in which all 10 bursts of EOD pulses were identical. We also constructed positive controls known to exhibit reliable responses, using 8 bursts of identical EODs (background), with the 9^th^ burst consisting of a novel waveform (a 90 degree phase-shifted EOD used during the first 8 bursts), followed by a 10^th^ burst of the original background waveform. Our experimental stimuli consisted of presenting eight bursts of the same *P. kingsleyae* EOD with the 9^th^ burst consisting of a different *P. kingsleyae* EOD. We computed the EOD rate of the subject fish by converting the EOD times of occurrence into a series of a delta functions, and convolving these a 300-ms Gaussian waveform (Carlson and Hopkins 2004). The specimen’s response was then recorded as the maximum EOD rate within a 2 sec window following each burst of EODs (Carlson and Arnegard 2011). The magnitude of the specimen’s response declined over repeated presentation of the background bursts (habituation). Therefore, we measured each specimen’s ‘novelty response’ as the change in EOD discharge rate (dishabituation) following the 9^th^ (novel) burst, relative to the EOD discharge rate following the 8^th^ presentation of the background burst. Using a Dunnett’s Test with Control (Dunnett 1955), we compared novelty responses toward a positive control stimulus and negative control. Significantly different novelty responses to experimental trains vs. negative control were considered evidence for discrimination between background and novel waveforms.

Positive control novel EOD stimuli were constructed by phase-shifting the background EOD waveform by 90°, which is maximally divergent in the time domain while preserving the frequency-spectrum. Phase shifting of EODs was accomplished by performing an FFT on the EOD waveform, followed by adding 90 degrees to the phase angle at each positive frequency, and subtracting 90 degrees from each the phase angle at each negative frequency. This was followed, in turn, by inverse FFT, which yielded a reconstructed EOD characterized by an altered (i.e., 90° shifted) phase spectrum with an unaltered power spectrum (Heiligenberg and Altes 1978; Hopkins and Bass 1981; Carlson and Arnegard 2011).

Some trials included ‘hybrid’ waveforms generated artificially from two normalized natural EOD waveforms: one P0 absent waveform from Bambomo and one P0 present waveform from Mouvanga. We centered these waveforms using the midpoint between peaks P1 and P2 of the waveform, and generated hybrids of varying P0 character by taking different weighted averages of the two waveforms (Fig. S1).

One set of experiments was performed within 6 hours of capture on P0-absent *P. kingsleyae* specimens from Bambomo creek (n=10), where both signal types are known to co-occur, as well as on P0-present *P. kingsleyae* individuals from Mouvanga creek (n=10), where only P0-present *P. kingsleyae* individuals occur but other *Paramormyrops* species are found. A second set of experiments was performed on additional P0-absent individuals from Bambomo Creek, which were captured as juveniles and transported to the Hopkins laboratory at Cornell University (Ithaca, NY), and reared to adulthood for further study (n=13). These fish were housed in community tanks with other individuals captured from their home stream. In addition, the fish were fed live blackworms daily, and they were maintained on a 12-hour light/dark cycle, with water temperatures maintained between 25-27°C, pH between 6.5-7.0, and water conductivity between 200-400µS.

### Electric organ histology & analysis

Serial sections of electric organs from selected individuals were made for light-microscopy analysis following the methods described by Gallant et al. (2011). Briefly, electric organs were removed from fixed specimens, decalcified overnight, dehydrated in a graded alcohol series, and infiltrated in glycol methacrylate resin (JB-4 resin; Polysciences, Inc). Serial parasagittal sections, each 6 µm thick, were cut from lateral to medial with a tungsten carbide microtome knife, mounted on glass slides, and stained with 0.5% Toluidine blue for 30 sec. For each specimen, we reconstructed one of four columns of electrocytes from these serial sections. Because each column of electrocytes surrounds the spinal cord, we began our reconstruction at the lateral edge of the electric organ, and stopped the reconstruction when the spinal cord was clearly visible (approximately 234-648 µm depending on the size of the individual). For each section, the number of stalk penetrations through each electrocyte was counted in, and averaged across, 50-70 electrocytes per section. An electrocyte was scored with a penetration whenever a stalk was observed to pass through either or both faces of the electrocyte. For our analysis, we considered each 6 µm section to have an independent number of penetrations from all other sections to minimize the probability of underestimating the total number of penetrations.

## Results

### Patterns of EOD variation

We assessed variation in EOD signals using both Principal Components Analysis (PCA; Fig. 4) and multidimensional scaling of signal cross correlations (MDS; Fig. S2) on each of the 327 individual EODs in this study.

**Figure 4.**
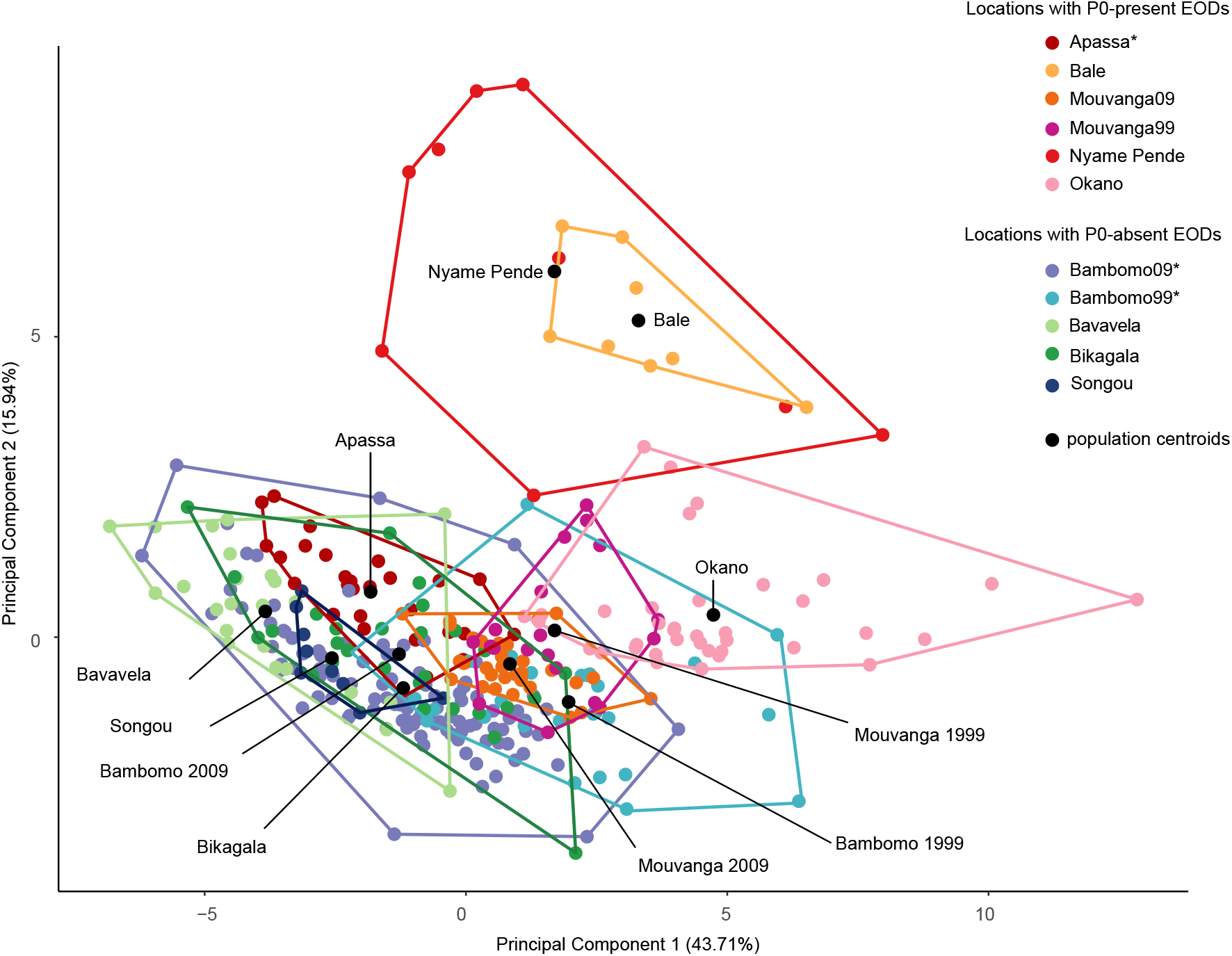
Principal component analysis of EOD waveform variation in 327 *P. kingsleyae* individuals from 11 populations in Gabon. Variation in waveforms was quantified using Principal Components Analysis (see Gallant et al. (2011) and text for further details). Variables related primarily to EOD duration loaded most strongly on principal component 1, whereas variables related to the magnitude of P0 loaded most strongly on principal component 2 (Table S3). Polygons enclose EOD waveforms from each recording locality. Polygon centroids are represented with black dots. Asterisks in the legend represent two populations with mixed signal types (Apassa is mostly composed of P0-present individuals, whereas Bambomo is mostly composed of P0-absent individuals).

The first principal component of our PCA related strongly to duration and explained 43.71% of the variation between individuals, while the second factor related strongly to the magnitude of P0 and comprised 15.94% of the variation between electric signals. Factor loadings for these two principal components are summarized in Table S3. In our MDS analysis, the number of dimensions was set to N = 2, which resulted in a stress of 0.0526, considered to give a good ordination representation of the cross-correlation matrix with low probability of misinterpretation (Clarke 1993).

Both MDS and PCA identified significant variation among recording localities (PERMANOVA on PCA signal distances: F_10,326_ = 58.62, *p*<0.001, PERMANOVA on MDS signal distances: F_10,326_ = 45.44, *p*<0.001). Mantel tests showed a strong and significant correlation between signal distances estimated from PCA and MDS (Fig. S3, Mantel test, *R* = 0.677, *p* = 0.001), which is also evident from comparison of Fig. 4 and Fig. S2. Interestingly, inter-population distances between Southern populations and Bale as well as between Southern populations and Nyame Pendé were larger with PCA-derived methods than with MDS-derived methods (Fig. S3). This is likely due to the fact that the PCA analysis included three variables pertaining to the frequency content of the EODs (peak frequency, low and high frequency with 3dB below the peak frequency) in addition to variables pertaining to the temporal content of the EODs. The cross-correlation analysis that the MDS was applied to, on the other hand, involves the progressive sliding of one waveform past the other, and thus indicates EOD similarity essentially in temporal content. Although a change in frequency content will be associated with a change in temporal content, the MDS analysis likely attributed less weight to specific frequency content differences. Regarding the two populations that were sampled in 1999 and again in 2009, we note that the overlap between population polygons corresponding to different sampling years is more extensive for Mouvanga than for Bambomo in both PCA and MDS, suggesting more extensive EOD divergence in Bambomo than in Mouvanga within a period of 10 years.

### Pattern of genetic variation

We were able to amplify fragments without any failed reactions (i.e. possible null homozyogotes) for all 327 individuals genotyped in this study. Total number of alleles detected at each of 5 loci over all populations ranged from 2-21 (Table 2). For each population, locus-specific expected heterozygosities ranged from 0.05-0.92 (Table 2). Exact tests produced no evidence of linked loci across all populations (p > 0.17). Of 55 unique locus-by-population combinations, only a few cases exhibited evidence of a deviation of observed heterozygosity from the expectation under Hardy-Weinberg equilibrium (at uncorrected p=0.05, see Table 2). After Bonferroni adjustment for multiple comparisons, only the NBB004 samples for Bambomo 1999, Bambomo 2009, and Nyamé-Pendé remain significant.

**Table 2.** Summary of loci attributes.

| *Pop.* | NBB01 |  |  |  | NBB02 |  |  |  | NBB03 |  |  |  | NBB04 |  |  |  | NBB05 |  |  |  |
| --- | --- | --- | --- | --- | --- | --- | --- | --- | --- | --- | --- | --- | --- | --- | --- | --- | --- | --- | --- | --- |
|  | *k* | *Ho* | *He* | *p* | *k* | *Ho* | *He* | *p* | *k* | *Ho* | *He* | *p* | *k* | *Ho* | *He* | *p* | *k* | *Ho* | *He* | *p* |
| Apassa | 6 | 0.71 | 0.67 | 0.955 | 4 | 0.79 | 0.71 | 0.393 | 6 | 0.75 | 0.68 | 0.212 | 4 | 0.64 | 0.75 | 0.276 | 17 | 0.75 | 0.92 | 0.184 |
| Bale | 4 | 0.50 | 0.53 | 0.655 | 8 | 0.75 | 0.89 | 0.432 | 7 | 0.75 | 0.85 | 0.762 | 3 | 0.50 | 0.49 | 0.283 | 9 | 0.75 | 0.92 | 0.241 |
| Bambomo 09 | 6 | 0.37 | 0.35 | 0.953 | 9 | 0.82 | 0.73 | 0.135 | 7 | 0.68 | 0.68 | 0.259 | 10 | 0.38 | 0.62 | 0.000 | 21 | 0.72 | 0.79 | 0.183 |
| Bavavela | 6 | 0.67 | 0.60 | 0.693 | 4 | 0.58 | 0.59 | 0.184 | 6 | 0.83 | 0.81 | 0.546 | 4 | 0.83 | 0.73 | 0.059 | 6 | 0.71 | 0.71 | 0.584 |
| Bambomo 99 | 4 | 0.41 | 0.38 | 0.537 | 7 | 0.67 | 0.77 | 0.251 | 6 | 0.44 | 0.64 | 0.006 | 8 | 0.22 | 0.69 | 0.000 | 13 | 0.78 | 0.81 | 0.455 |
| Bikagala | 5 | 0.68 | 0.64 | 0.485 | 4 | 0.50 | 0.50 | 0.04 | 4 | 0.50 | 0.59 | 0.437 | 4 | 0.34 | 0.33 | 0.373 | 4 | 0.57 | 0.54 | 0.053 |
| Mouvanga 09 | 2 | 0.09 | 0.09 | 1.000 | 3 | 0.16 | 0.15 | 1.000 | 4 | 0.25 | 0.23 | 1.000 | 6 | 0.44 | 0.43 | 0.445 | 3 | 0.44 | 0.51 | 0.576 |
| Mouvanga 99 | 2 | 0.05 | 0.05 | 1.000 | 3 | 0.16 | 0.15 | 1.000 | 5 | 0.26 | 0.41 | 0.023 | 6 | 0.37 | 0.48 | 0.228 | 4 | 0.63 | 0.56 | 0.644 |
| Nyamé Pendé | 2 | 0.11 | 0.11 | 1.000 | 10 | 0.67 | 0.91 | 0.046 | 7 | 1.00 | 0.81 | 0.691 | 2 | 0.00 | 0.47 | 0.004 | 10 | 0.89 | 0.88 | 0.993 |
| Okano | 7 | 0.53 | 0.55 | 0.271 | 2 | 0.24 | 0.25 | 0.569 | 7 | 0.42 | 0.37 | 1.000 | 8 | 0.45 | 0.43 | 0.489 | 16 | 0.82 | 0.90 | 0.197 |
| Songou | 3 | 0.22 | 0.57 | 0.014 | 2 | 0.11 | 0.11 | 1.000 | 4 | 0.33 | 0.65 | 0.12 | 2 | 0.22 | 0.52 | 0.172 | 2 | 0.22 | 0.21 | 1.000 |

For each locus and population, the number of alleles (*k*) at each locus, the observed (*H<sub>o</sub>*) and expected heterozygosity (*H<sub>e</sub>*) are reported. Probabilities of deviation from HWE are reported as *p*

The allele frequency histograms at each of 5 loci for each of the 327 individuals in the full dataset are shown in Figure 5. For the two populations that were sampled in 1999 and again in 2009 (Mouvanga and Bambomo), allele frequencies appeared to change little over this ten year period between sampling. Between populations, there are apparent differences between the distribution of alleles; of particular note are the presence of several alleles (NBB002, ∼250 bp, NBB005 several alleles between 300-400 bp), which are present in Bambomo and Apassa creeks but absent in other populations.

**Figure 5.**
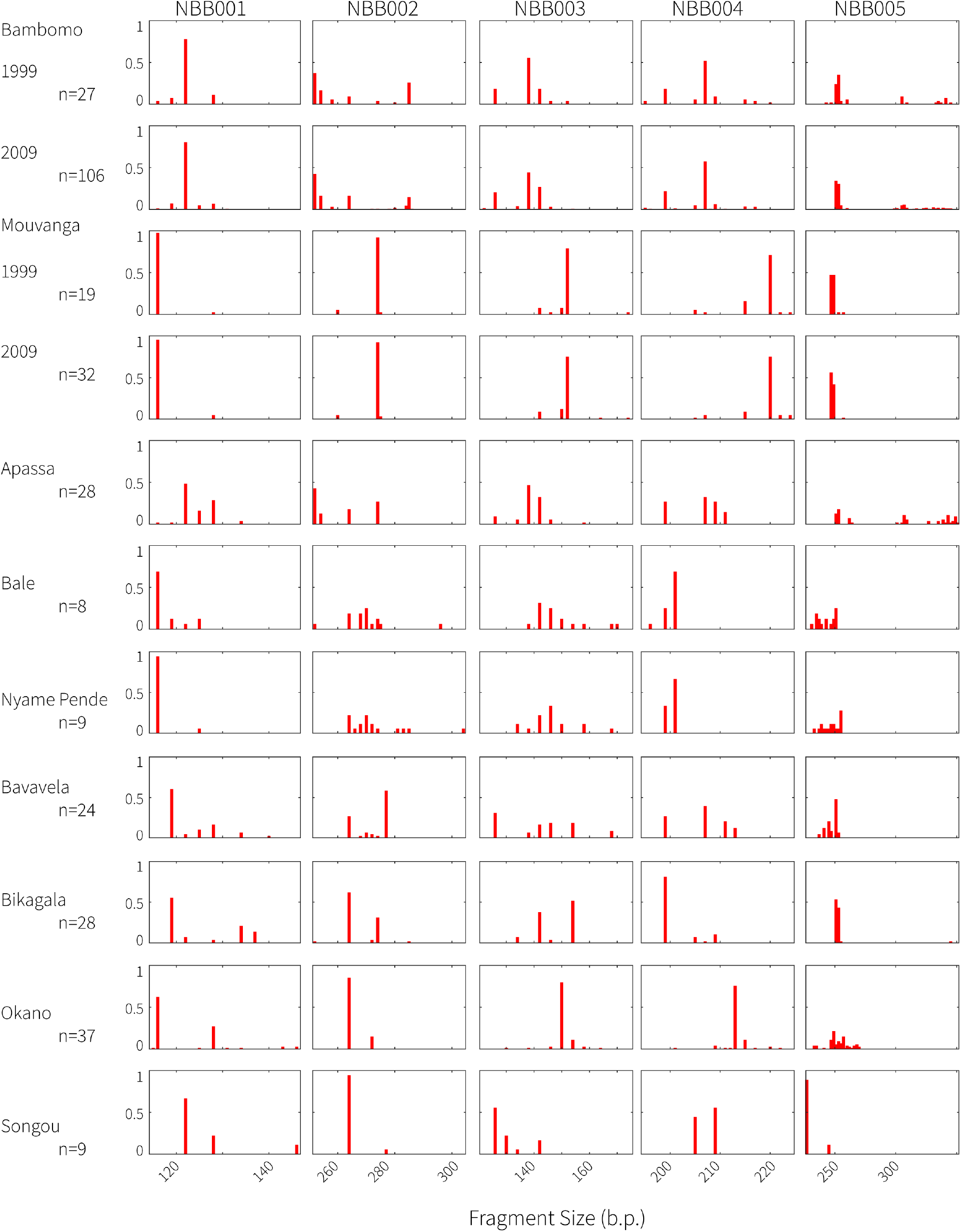
Allele frequency histograms for each population and microsatellite locus. Microsatellite loci were designated as NBB001-NBB005 by Arnegard et al. (2005). For each population, sample sizes are reported as number of individuals genotyped, so the plots include twice as many allele copies.

F_st_ values of all pairwise comparisons between the *P. kingsleyae* populations surveyed are shown in Figure 6. All populations were significantly differentiated from one another at the p=0.05 and the Bonferonni-corrected threshold (p=0.001), with the exceptional pairwise comparisons of Bambomo Creek 1999 vs. 2009, Mouvanga Creek 1999 vs. 2009, and Nyamé-Pendé vs. Bale Creek. Among all pairs of populations, the magnitude of genetic differentiation between populations varied from F_st_ = 0.055 to F_st_ = 0.65. We note that Mouvanga was highly differentiated from all other populations, with F_st_ values ranging from 0.377 (Mouvanga99-Nyame Pende) to 0.65 (Mouvanga09-Songou). Most importantly, the populations of Bambomo (predominantly biphasic individuals) and Apassa (predominantly triphasic individuals), which represent a potential hybrid zone, exhibit the lowest F_st_ values of our dataset (Fst<0.06, Fig. 5), despite being phenotypically distinct with respect to P0-presence/absence.

**Figure 6.**
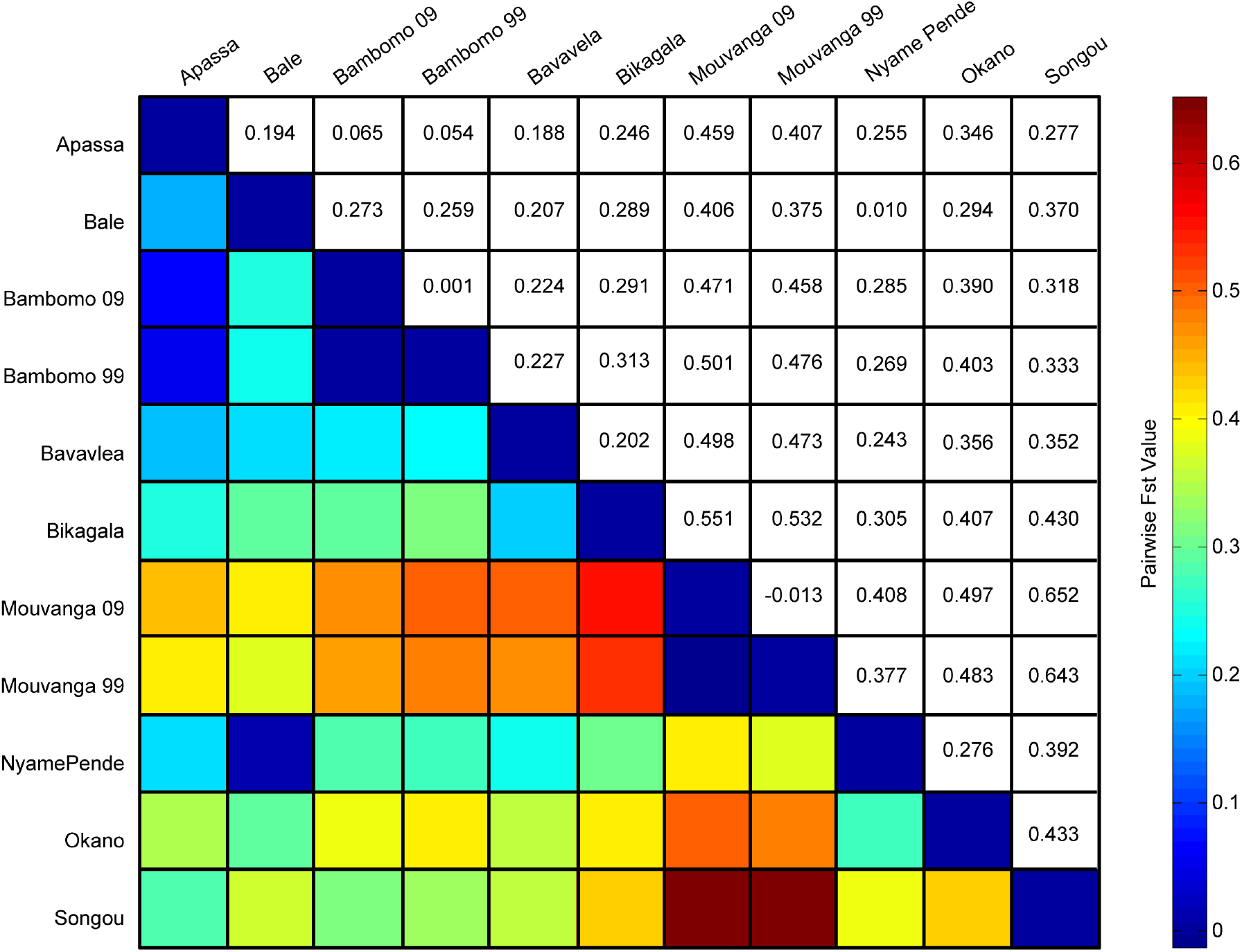
F_st_ values for each pairwise comparison of populations. Lower half of the matrix codes F_st_ value by color, the upper half of the matrix reports the individual F_st_ values. All pairwise comparisons of F_st_ values were significant at Bonferroni-corrected thresholds, with the exception of Mouvanga Creek (1999 vs. 2009), Bambomo Creek (1999 vs. 2009), and Bale Creek vs. Nyame Pende Creek.

### Isolation by distance

We investigated the relationship between genetic differentiation and geographic distance, measured as shortest river distance between populations (Fig.7A). Among all population pairs in Gabon, we found no significant relationship between genetic and geographic distance (Mantel test, *R*=-0.042, *p*=0.568), even after correcting for potential regional-level effects (partial Mantel test *R*=0.099, p=0.278). These results were consistent across different genetic distance measures (Table S4) or when samples collected in different years in Bambomo and Mouvanga were treated as one instead of separate populations (Table S5&S6).

**Figure 7.**
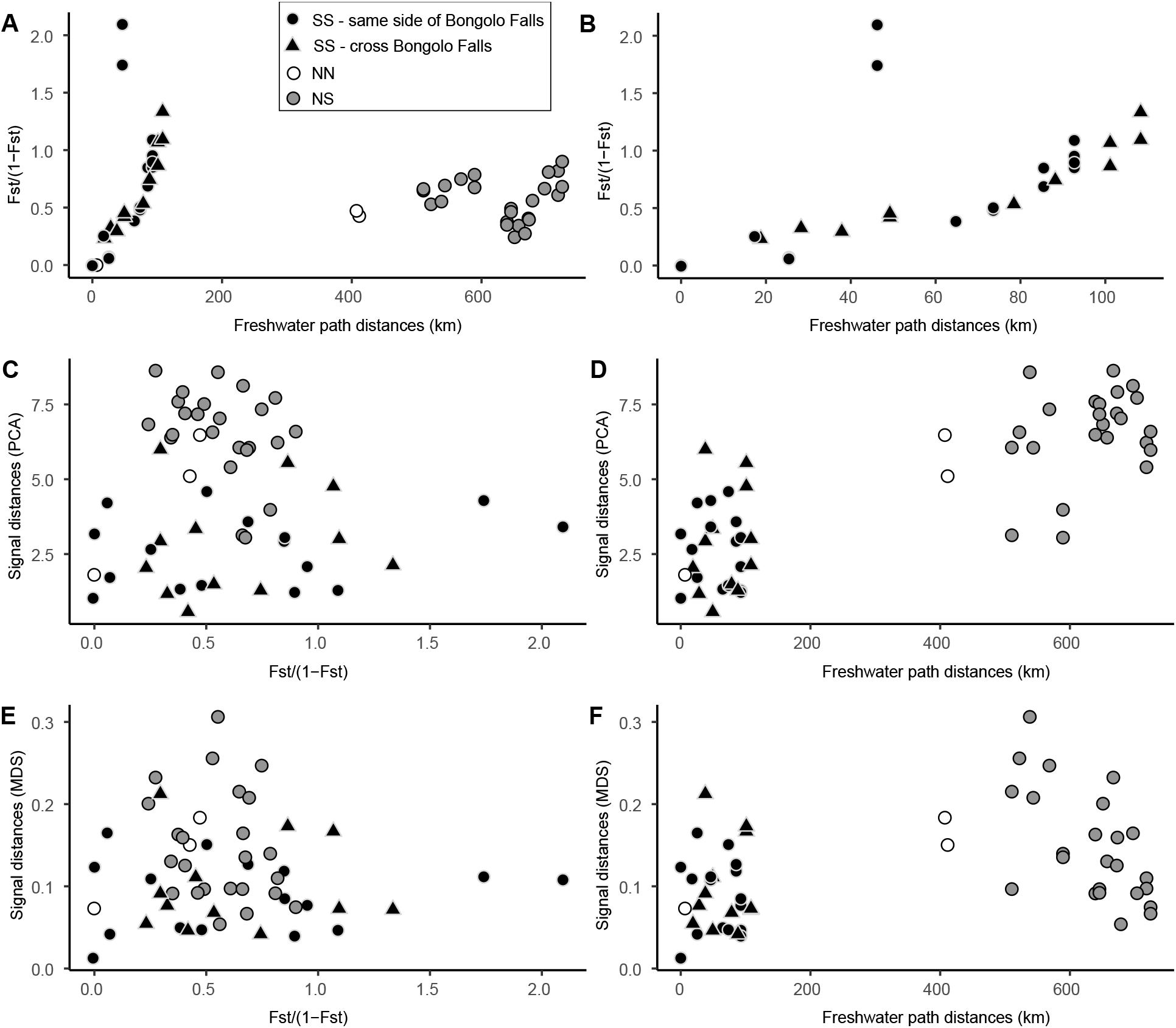
Correlation plots between genetic, signal and geographic distances for all populations (n=11populations, all symbols) and within Southern populations only (n=8 populations, black symbols). A. Population pairwise genetic distance in relation to geographic distance. **B.** Pairwise genetic distance in relation to geographic distance in Southern populations only. **C.** Correlation between PCA-derived signal and genetic distances. **D.** Correlation between PCA-derived signal and geographic distances. **E**. Correlation between MDS-derived signal and genetic distances. **F.** Correlation between MDS-derived signal and genetic distances. (SS: South-South comparisons, NN: North-North comparisons, NS: North-South comparisons).

In contrast, among population pairs restricted to the South of Gabon, genetic distance was strongly related to geographic distance (Fig. 7B; R= 0.576, *p =* 0.02), regardless of the genetic distance metric used (Table S4) or whether Bambomo and Mouvanga samples from different years were treated as independent populations or not (Table S5&S6). However, we found that genetic distances between populations separated by Bongolo falls were on average no greater than those between populations on the same side of the Falls (Fig. S4). This was confirmed by the lack of correlation between genetic distance and the Bongolo model matrix (Table 3).

**Table 3.**
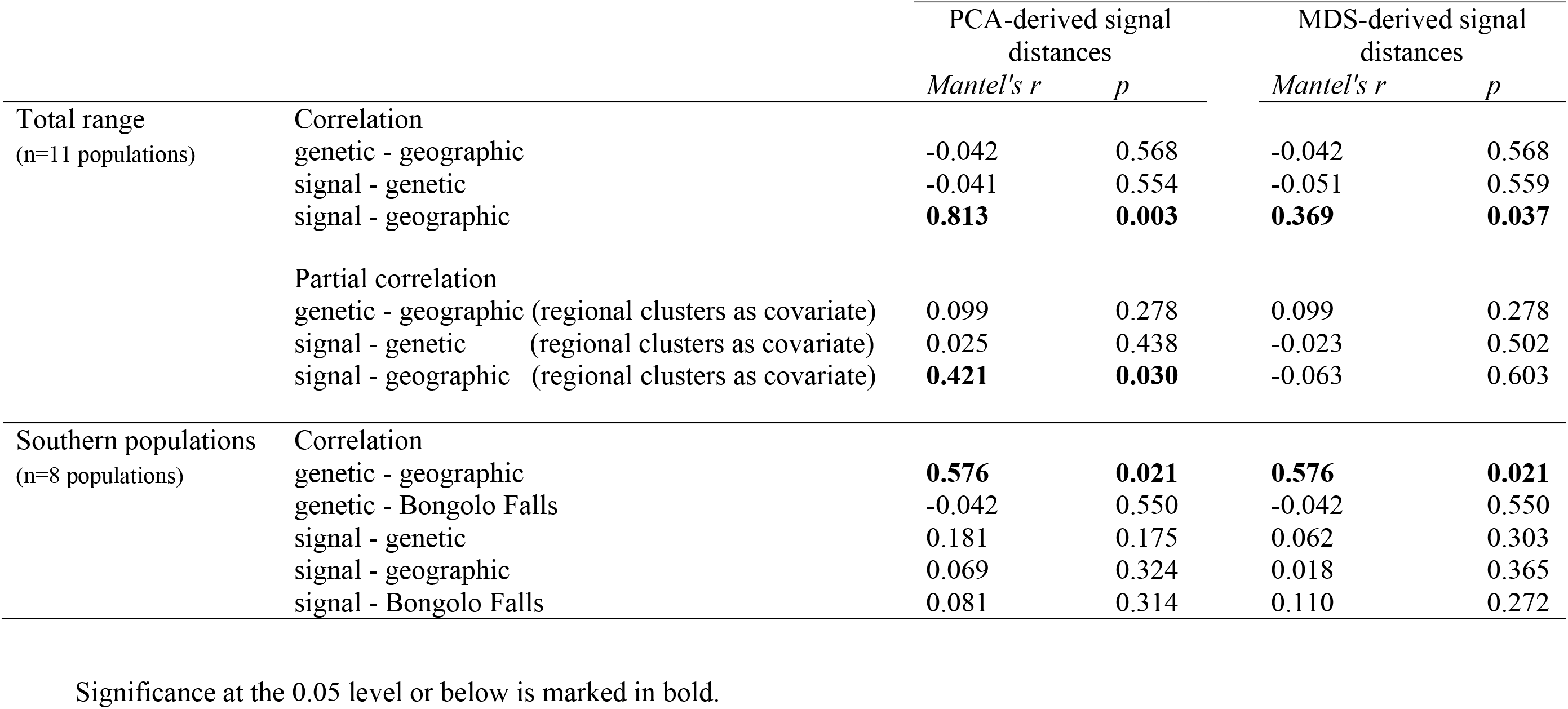
Results of standard and partial Mantel tests between genetic distances (Fst/(1-Fst)), signal distances (obtained through PCA (left) and MDS (right)), geographic distances (km), and presence/absence of Bongolo Falls (Southern populations) at two spatial scales.

### Correlation between signal, genetic, and geographic distances

We investigated the relationship between signal, genetic and geographic distance (Fig. 7 C,D,E,F) at two spatial scales (for correlation plots focusing only on Southern population, see Fig. S5 A,B,C,D). When considering the entire dataset, signal distances were not correlated with genetic distances (Fig.7 C&D, Table 3), even after correcting for potential regional-level effects (partial Mantel test *R*=0.025 p=0.438). These results were consistent regardless of how signal or genetic distances were estimated (Table 3, Table S4) and of whether samples collected from different years in Bambomo and Mouvanga were considered as one instead of separate populations (Table S5&S6).

On the other hand, we found a strong and significant relationship between signal and geographic distance using PCA-derived signal distances (Fig. 7D, Table3), even after correcting for potential regional-level effects (partial Mantel test *R*=0.421 p=0.03) and rafter treating samples collected from different years as one instead of separate populations (TableS5&S6). Using MDS-derived signal distances, the signal-geography correlation was weaker yet significant (Fig. 7F, Table 3); however, it disappeared when we controlled for regional-level effects (partial Mantel test=-0.063, p=0.603). The same results were found regardless of how Bambomo and Mouvanga samples were grouped (Table S5,S6).

When considering only Southern populations, none of the variables tested (genetic distance, geographic distance, and Bongolo Falls model matrix) were found to correlate with signal distances (Fig. S5A,B,C,D, Table 3, Table S4,S5,S6).

### P. kingsleyae can detect intraspecific EOD waveform variation

We utilized a habituation-dishabituation paradigm (Carlson et al. 2011) to determine whether individual *P. kingsleyae* were capable of discriminating between sympatric and allopatric *P. kingsleyae* EOD waveforms that were either P0-absent or P0-present (Fig. 8A,B). We performed two sets of experiments to assess this: the first set of experiments was performed with single stimulus presentations and recordings of responses across 20 field-captured individuals (Fig. 8C,D), whereas the second was performed with 5 repetitions of randomly interleaved stimulus trains to laboratory maintained individuals, which enabled us to average responses and thereby reduce variability (Fig9E-H; see Methods).

**Figure 8.**
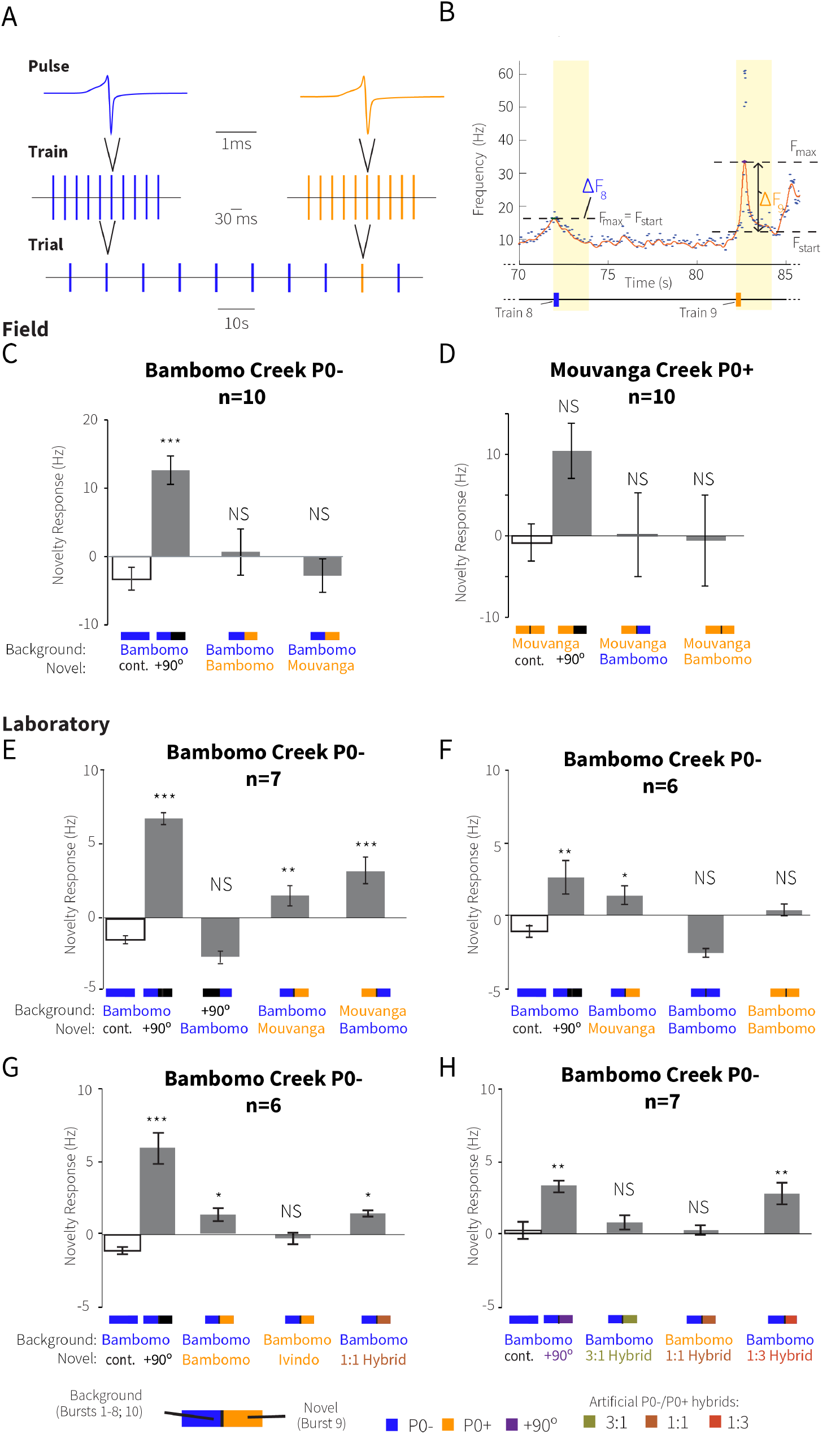
Results of dishabituation playback. Playback experiments performed on *P. kingsleyae* **A**. Stimuli used: 10 trains of 10 EODs each are presented in each trial. 8 trains were comprised of one repeated EOD waveform type (background) followed by one train of 10 EODs (novel) followed by one train of 10 background EODs. **B.** Example of subject response. Subject’s EOD discharges were continuously monitored (black dots), and discharge frequency was fitted with a spike density function (red line). The maximum and starting discharge rate of the SDF was calculated over the 2sec interval from the onset of each train presentation. The difference between these values defined the change in frequency (ΔF) for that train. Novelty response was defined as the difference in ΔF between the 8^th^ and 9^th^ (novel) train presentation. See methods for details. **C-D** Experiments performed on field captured specimens. **E-H** Experiments performed in the laboratory. Subjects and sample sizes are described in italics above each plot. Bars show mean novelty responses averaged for all specimens. Subtext indicates the localities where background and novel waveforms were recorded, if applicable. Stars reflect significance as compared to negative control (Dunnett’s Test w/Control: NS: not significant, * p=0.05; ** p<0.01; *** p<0.001.

**Figure 9.**
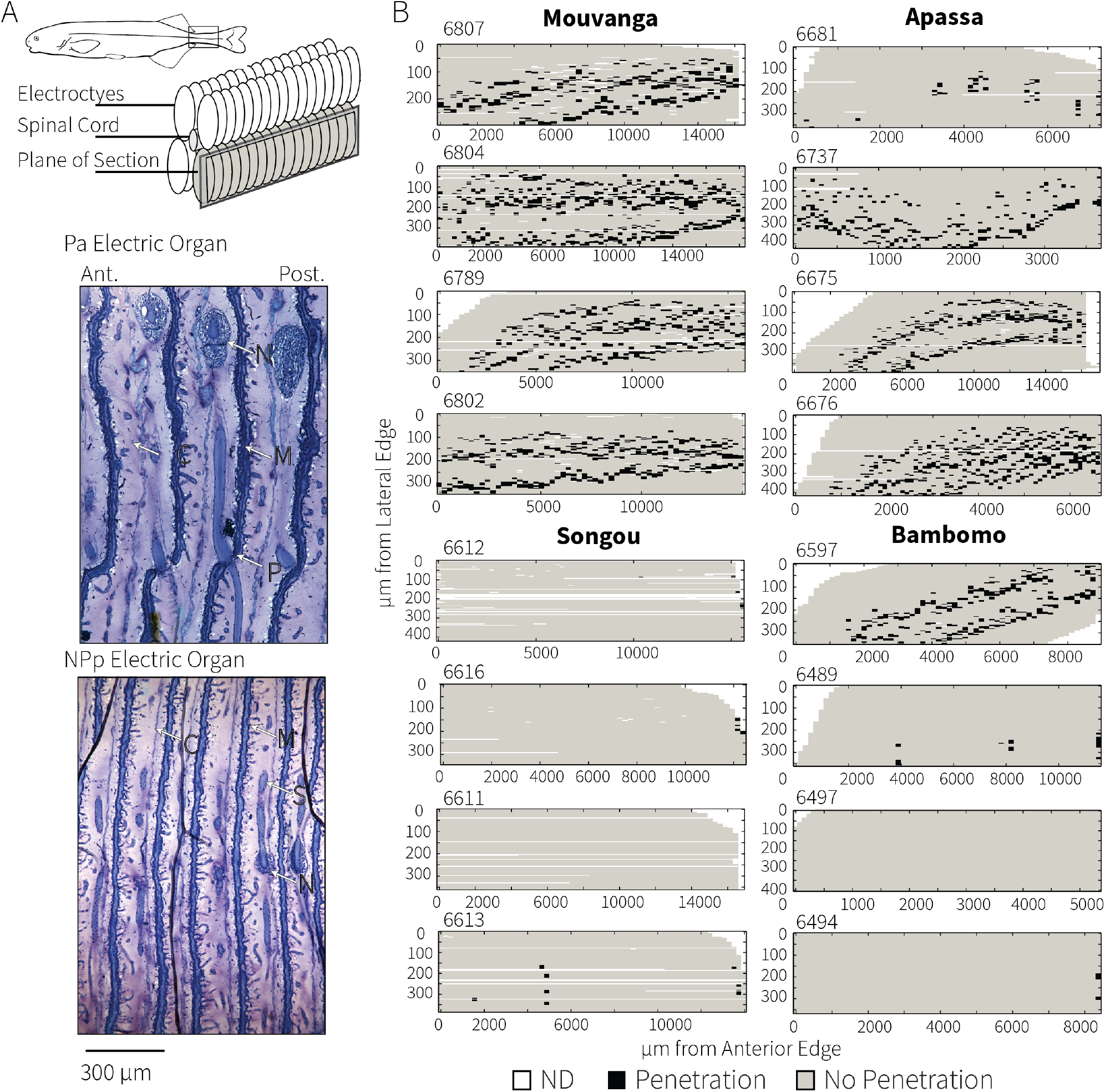
Summary of histological survey of electric organs. **A.** Individual *P. kingsleyae* were sectioned sagittally from lateral to medial, following one column of electrocytes of the 4 that comprise the electric organ. The electric organ is restricted to the caudal peduncle in mormyrids. Example histology showing the two basic types of anatomical configuration in the electric organ *Pa* (penetrating with anterior innervation) and NPp (nonpenetrating with posterior innervation). Stalks (S) can be seen clearly passing through the electrocyte penetrations (P) in Pa but not in NPp type electrocytes. Microstalklets (M) can be observed on the posterior face in both cases, and connective tissue boundaries (C) bounding each electrocyte are indicated. Innervation (N) can be observed on the anterior or posterior side of the electrocyte accordingly. Pa type electrocytes result in EOD signals that have a small head negative phase P0, which is absent in individuals with electrocytes that are NPp (see Gallant et al., 2011). **B.** Examples of histological analysis of 4 electric organs from each population. Each pixel represents an individual electrocyte in an individual section that was scored for presence of penetrations (black). In Apassa creek, most individuals have Pa type observed in each electrocyte from anterior to posterior, whereas one individual had NPp type and Pa type electrocytes (patches of black in mostly grey background). In Bambomo, most individuals were NPp with one individual exhibiting all Pa type electrocytes. Songou was entirely comprised of individuals with NPp type electrocytes, and Mouvanga was comprised of individuals with entirely Pa type electrocytes. Full analysis is summarized in Table 4.

In the first field experiment (Fig. 8C), P0-absent *P. kingsleyae* from Bambomo creek showed behavioral evidence of discriminating phase-shifted EODs from P0-absent EODs (p<0.001), but did not show evidence of discriminating between P0-absent EODs (background) and Mouvanga P0-present EODs or Bambomo P0-present EODs (novel). In the second field experiment (Fig. 8D), P0-present *P. kingsleyae* from Mouvanga creek showed greater dishabituation to phase shifted EODs than test EODs, but this difference was not significant (p=0.1751). There was no significant difference in response between Mouvanga P0-present EODs (background) and Bambomo P0-absent EODs or Bambomo P0-present EODs (novel). Taken together, these field experiments performed on *P. kingsleyae* from Bambomo Creek and Mouvanga creek did not support the hypothesis that *P. kingsleyae* are able to discriminate between the subtly different P0-absent and P0-present waveforms. We therefore performed laboratory experiments that were designed to be more sensitive at detecting small differences in response through averaging.

In the first laboratory experiment (Fig. 8E), we tested the hypothesis that *P. kingsleyae* from Bambomo Creek are able to discriminate allopatric P0-present EODs from sympatric P0-absent EODs. We were able to determine a statistically significant response to the positive control (novel=90° phase shifted EOD; p<0.001). In addition, we detected statically significant changes in novelty responses to stimulus trains where both Mouvanga P0-present EODs were novel (p<0.01), and where Bambomo P0-absent EODs were novel (p<0.001). This evidence is consistent with the hypothesis that *P. kingsleyae* are able to discriminate between P0-present and P0-absent waveforms, but is confounded by the fact that allopatric populations, and even individuals, can differ slightly in EOD waveform (Gallant et al. 2011). Thus, there may be cues other than P0 that mediated this discrimination.

To rule out the possibility of individual EOD discrimination on the basis of characters *other than* P0-presence/absence, we tested *P. kingsleyae* captured in Bambomo Creek with individual sympatric EOD waveforms where P0 <u>was not</u> available as a cue (Fig. 8F). Though subjects responded significantly to positive controls (+90° and Mouvanga which have large P0-absent waveforms; p<0.001), subjects did not exhibit a significant novelty response when presented background and novel stimuli that were all P0-absent (p=0. 3478), or when background and novel stimuli were all P0-present (p=0. 3869). In a third, corollary laboratory experiment, we tested the hypothesis that *P. kingsleyae* from Bambomo Creek are able to discriminate between sympatrically occurring EOD variants (Fig. 8G) where P0-presence/absence <u>was</u> available as a cue. Subjects responded significantly to positive controls (p<0.001), and to P0-present waveforms from Bambomo in a background of P0-absent waveforms from Bambomo (p=0.02), though not to P0-present waveforms from Ivindo (which have very large P0s; p=0. 7189) from a background of P0-absent waveforms. Together, these three laboratory experiments support the hypothesis that *P. kingsleyae* can differentiate between sympatric EOD waveforms when P0 is available as a cue.

Finally, in the fourth laboratory experiment (Fig. 8H), we examined the ability of *P. kingsleyae* to discriminate between artificial EOD waveforms that had very small P0 peaks present (1:1 P0-absent/P0-present hybrid; see methods), and natural EOD waveforms that had no P0-present. We determined a statistically significant response to positive controls (p<0.01), and to EODs that represented 1:3 ratios of P0-absent to P0-present waveforms (p<0.01), however, subjects did not elicit a statistically significant response to 3:1 (p=0.873) or 1:1 ratios (p=1.0) of P0-absent to P0-present EODs.

The results of the field and laboratory experiments are consistent with the hypothesis that *P. kingsleyae* can differentiate between P0-absent and P0-present waveforms.

### Morphological analysis of electric organs identifies additional specimens with mixed anatomy in Apassa creek

Gallant et al (2011) showed that P0-absent EODs are produced by fish with *NPp* anatomy, while P0-present EODs were produced by fish with penetrating stalked electrocytes in the electric organ. In addition, Gallant et al (2011) presented evidence for electric organs with mixed morphology, wherein some electrocytes had penetrating stalks while other electrocytes from the same organ had non-penetrating stalks. We confirmed this observation with an additional analysis of 21 electric organs collected in 2009. Our present analysis supports the existence of these morphological types. In Fig. 9 we show 4 examples of electric organs surveyed in Bambomo, Apassa, Mouvanga, and Songou Creeks, and summarize our analysis of all 21 individuals in Table 7. In Apassa and Mouvanga all individuals were of the Pa morphology type, with one exceptional individual in Apassa exhibiting mixed morphology (*NPp* + *Pa* morphology in the same individual). In Bambomo Creek and Songou creek, we detected individuals that had entirely *NPp* type morphology, with one exceptional individual in Bambomo creek that had entirely *Pa* type morphology.

**Table 4.** Summary of electric organ histological survey, performed on 21 specimens of *Paramormyrops kingsleyae*. For each individual, the collection locality and specimen number are provided. For each specimen, one column of the entire electric organ was surveyed for the specified depth (µm) from lateral to medial (see Fig. 8). Total number of electrocytes, and the number of those exhibiting *Pa* type morphology and *NPp* type morphology are provided. Finally, the assessment of the overall EO morphology is specified as either Pa (>75% of electrocytes Pa) NPP (>75% electrocytes NPp) or mixed (<75% of electrocytes of one type). Note that some individuals determined to have Pa anatomy did not have penetrations in the rostral or caudal portion of the organ (See Fig. 8). In these cases, electrocytes could not be fully surveyed because of the orientation of tissue during sectioning.

| Population | Specimen No. | µm Surveyed | No. Sections | No. Electrocytes | No. *Pa* | No. *NPp* | Morphology |
| --- | --- | --- | --- | --- | --- | --- | --- |
| Apassa | 6681 | 420 | 70 | 60 | 15 | 45 | Mixed |
| Apassa | 6675 | 648 | 108 | 64 | 50 | 14 | Mixed |
| Apassa | 6676 | 366 | 61 | 69 | 58 | 11 | Pa |
| Apassa | 6637 | 258 | 43 | 70 | 65 | 5 | Pa |
| Bambomo | 6489 | 240 | 40 | 60 | 10 | 50 | NPp |
| Bambomo | 6494 | 384 | 64 | 61 | 1 | 60 | NPp |
| Bambomo | 6497 | 276 | 46 | 67 | 0 | 67 | NPp |
| Bambomo | 6500 | 270 | 45 | 59 | 1 | 58 | NPp |
| Bambomo | 6547 | 378 | 63 | 52 | 1 | 51 | NPp |
| Bambomo | 6549 | 186 | 31 | 60 | 5 | 55 | NPp |
| Bambomo | 6597 | 438 | 73 | 58 | 43 | 15 | Mixed |
| Mouvanga | 6789 | 516 | 86 | 58 | 53 | 5 | Pa |
| Mouvanga | 6811 | 342 | 57 | 62 | 55 | 7 | Pa |
| Mouvanga | 6807 | 582 | 97 | 50 | 49 | 1 | Pa |
| Mouvanga | 6810 | 390 | 65 | 66 | 61 | 5 | Pa |
| Mouvanga | 6804 | 570 | 95 | 65 | 65 | 0 | Pa |
| Mouvanga | 6802 | 570 | 95 | 57 | 57 | 0 | Pa |
| Songou | 6611 | 252 | 42 | 59 | 0 | 59 | NPp |
| Songou | 6612 | 564 | 94 | 70 | 4 | 66 | NPp |
| Songou | 6613 | 234 | 39 | 64 | 5 | 59 | NPp |
| Songou | 6616 | 480 | 80 | 57 | 2 | 55 | NPp |

## Discussion

In this study, we tested the overall hypothesis that genetic drift is responsible for the biogeographic distribution of EOD signals in *Paramormyrops kingsleyae*. We found no significant relationship between signal and genetic distance, implying that electric signal divergence in *P. kingsleyae* cannot be explained by drift alone. Next, we tested the ability of individuals collected from this hybrid zone to discriminate divergent signal types and examined genetic and morphological data to investigate whether divergent EODs could be used as cues for assortative mating and thus rather be shaped by sexual selection. These experiments demonstrated that *P. kingsleyae* are able to discriminate between P0-present and P0-absent EODs and support the existence of a previously hypothesized ‘hybrid zone’ between triphasic and biphasic *P. kingsleyae*. Taken together, our findings show that *P. kingsleyae*’s signals do not diverge at a rate that is consistent with neutral evolution and that selective forces are more likely candidates driving EOD diversity in this species.

A key prediction of the drift hypothesis is that EOD signal variation should be highly correlated with variation in neutral genetic markers. However, our results, which encompass two independent measures of signal variation (i.e. PCA and MDS), demonstrate that phenotypic differentiation in *P. kingsleyae* does not correspond to genetic distance between populations at any scale of our study (Table 3). Although we detected significant levels of genetic differentiation between all populations separated by > 600 km, we note that genetic differentiation patterns were not consistent across all scales of geographic isolation in our study. In the South, we found a highly significant relationship between genetic differentiation and geographic distance in as would be predicted by isolation-by-distance models (Wright 1984). The presence of Bongolo Falls was, however, not found to explain any of the genetic structure in this region (Table 2, Table S3,S4,S5), implying that populations separated by this 15m waterfall are not necessarily genetically more diverged than populations not separated by it. We speculate that this may be caused by occasional downstream migration from populations living above the Falls. Interestingly, we also note that in Bambomo and Mouvanga Creeks, allele frequencies were remarkably stable over a 10 year period, suggesting large effective population sizes (Waples 1989). When considering all populations, we did not find a significant relationship between genetic and geographic distance, suggesting that patterns of genetic differentiation are due to other processes than isolation by distance. Since the biogeographic distributions of tropical freshwater fishes are mainly constrained by regional landscape and ecological features such as basin geomorphology and historical connections and disconnections between adjacent river basins, differentiation patterns in these taxa at large spatial scales are often purely the result of vicariance and drift (Albert and Reis 2011). The observed pattern in *P. kingsleyae*’s genetic structure across the entire country could be related to the balance between mutational processes within populations and gene flow between them (Hutchison and Templeton 1999), whereby gene flow and drift influence regional population structure differently depending on scale. As microsatellites have very high mutation rates and unique mutational processes (Selkoe and Toonen 2006), it is possible that between our isolated Southern and Northern regions, random mutations due to drift in different populations to the same alleles (i.e. homoplasy) reduced F_st_ estimates. Future phylogenetic and biogeographic studies using more numerous biallelic markers (i.e. single nucleotide polymorphisms) will be necessary to fully understand *P. kingsleyae’*s genetic structure and colonization history over the Gabonese landscape.

The lack of correlation between electric signal and genetic distance in *P. kingsleyae* contrasts with a previous study on the gymnotiform *Brachyhypopomus occidentalis*, the only other electric fish species for which the role of drift in signal divergence has been tested, which reported strong correlations between neutral genetic and signal distances (Picq et al. 2016). Although drift has been found to play an important role in the divergence of communication signals in several taxa including electric fish (Picq et al. 2016), Neotropical singing mice (Campbell et al. 2010), Amazonian frogs (Amézquita et al. 2009), and greenish warblers (Irwin et al. 2008), it is important to note that most studies testing the contribution of drift in communication signals rarely report evidence for neutral signal evolution (Soha et al. 2003; Nicholls et al. 2006; Pröhl et al. 2006; Ruegg et al. 2006; Pröhl et al. 2007; Rudh et al. 2007; Dingle et al. 2008; Huttunen et al. 2008; Tobias et al. 2010; Cadena et al. 2011). Selective forces therefore are often reported as the main contributors to communication signal divergence.

Electric signal differences were more likely to be predicted by geographic distance than by genetic distance at the scale of the whole country. The correlation between PCA-derived signal distances and geographic distance was high and significant even after correcting for potential regional-level effects, a result consistent with previous findings from Gallant et al. (2011). This correlation, however, did not hold when using MDS-derived signal distances and correcting for potential regional-level effects. This difference is most likely caused by the fact that landmarks pertaining to the EOD frequency content were explicitly included as part of the 21 total EOD measurements in the PCA analysis, and not in in the MDS analysis. Of the three frequency-content variables included in the PCA (Fig. 3, Table S2) only high frequency content (*ffthi,* frequency above the peak power frequency at −3dB) was found to be significantly different between Southern populations (all triphasic with large P0-magnitude) and Northern populations (triphasic with small P0-magnitude and biphasic populations), (mean *ffthi* for Southern populations = 2777.90 Hz, mean *ffthi* for Northern populations = 1585.11 Hz, t-test *t*=-14.96, *p*<0.001). Interestingly, low-capacitance objects such as small invertebrate larvae attenuate higher-frequencies more readily (Meyer 1982; von der Emde and Ringer 1992; Crampton 1998). It is therefore possible that EODs differing in high-frequency composition convey differential electrolocation capacities. This observation suggests that additional forces such as natural selection for electrolocating different prey items in different habitats may also be contributing to the evolution of EODs in *P. kingsleyae*. Future studies will have to investigate whether *P. kingsleyae* with different signal types actively prey on different items and also look at what abiotic or biotic factors are potentially correlated with this clinal variation in EOD frequency content across Gabon.

Our laboratory experiments provide evidence that *P. kingsleyae* from the putative hybrid zone are capable of discriminating between P0-present and P0-absent EODs (Fig. 8). This finding is consistent with several studies that have characterized the neural encoding of EOD waveforms (Amagai et al. 1998; Friedman and Hopkins 1998; Xu-Friedman and Hopkins 1999; Carlson 2009; Carlson and Arnegard 2011; Baker et al. 2013; Lyons-Warren et al. 2013). These results indicate that in zones of signal sympatry, the observed EOD variation has the potential to be meaningful to *P. kingsleyae* receivers and there could play a role in communication within this species.

Our genetic data pointed out to weak but significant genetic differentiation between Apassa and Bambomo (Fst<0.07), where both signal types occur in sympatry, despite the fact that these populations are nearly fixed for alternate electric organ anatomies and signal types. This suggests that these populations are able to maintain phenotypic divergence in the face of gene flow (i.e. hybridization). Given the large fluctuations in water level due to rainy season (Arnegard et al. 2006), it is plausible that these sites are periodically connected through seasonal flooding events, allowing migration and gene flow to occur between these otherwise geographically isolated populations. Another line of evidence supporting hybridization is that both populations exhibit a so-called “rare alleles” phenomenon (Fig. 5): both have additional alleles that are absent from all other populations genotyped, which has been demonstrated as a signature of hybrid zones in other taxa (Golding 1983; Barton 1985; Hoffman and Brown 1995). A third, and most compelling line of evidence for hybridization is the existence of mixed electric organ morphology individuals in both Apassa and Bambomo (Fig. 9 and Table 4 in addition to those already discovered in Gallant et al. (2011)). Specimens with both stalk geometries in electrocytes were obtained in hybrids of artificial crosses between the mormyrid *Campylomormyrus tshokwe* and *Campylomormyrus tamandua* (Kirschbaum et al. 2016), strongly suggesting that the same electrocyte pattern in *P. kingsleyae* does result from hybridization between different EOD morphs.

To summarize, we report evidence that *P. kingsleyae* are perceptually capable of detecting minute EOD differences characteristic of neighboring populations. However, considerable evidence of hybridization suggests the availability of this information to the nervous system has not led to complete reproductive isolation between EOD morphs. This could be partly explained by the considerable variation in discrimination ability between individuals that we found in our field and laboratory experiments, despite the overall evidence of discrimination (Fig. 8). Variation in discrimination and perceptual biases among choosers in the context of mate choice has been demonstrated extensively (Rodríguez and Andrew Snedden 2004; Ryan and Cummings 2013) and can have important implications for evolutionary diversification. It is therefore possible that the variation in discrimination response in our experiments accurately reflects that not all *P. kingsleyae* are equally good at discriminating EOD waveforms, which may be due to differences in reproductive state, age, condition and sensory acuity of the individuals (Rosenthal 2017). This discrimination variation could in turn lead to variation in the strength of preference for a given signal type among individuals, which could be a mechanism explaining the dual maintenance of different EOD types and the existence of a few hybrid individuals. We note that discrimination ability does not equate signal preference, although in order to have a preference, individuals must be able to perceive signal differences. In the same way, a lack of novelty response in our experiments does not necessarily mean lack of discrimination, as some fish may be able to distinguish signals and still not show a behavioral response due to different internal or external factors. These results motivate a more detailed study of waveform preferences in this species in the context of mate choice to determine the degree of assortative mating that may occur on the basis of EOD waveform. It remains possible that the observed variation in *P. kingsleyae* EODs is also important for other sender-receiver dynamics such as intraspecific competition or resource defense.

Overall, our study suggests that the hypothesized role of genetic drift (Gallant et al. 2011) is insufficient to explain the evolution of *P. kingsleyae* waveforms. We found that *P. kingsleyae* possess the perceptual ability to discriminate between P0-present and P0-absent EODs, but also found evidence of hybridization between EOD morphs, suggesting that reproductive isolation with respect to EOD type in *P. kingsleyae* is not complete. Field and laboratory playback experiments among *Paramormyrops* species have revealed strong preferences for species-specific EOD waveforms (Hopkins and Bass, 1981) during courtship, indicating that EODs are most likely involved in maintaining pre-zygotic isolation between closely related species. Our data suggest that divergence in EODs occurs at the population level and is unlikely to be driven by drift alone, thereby stimulating new avenues of research focusing on selective forces in order to understand the evolutionary processes shaping signal diversification.

## Author Contributions

SP performed signal and genetic variation analyses and co-designed the study. CC performed histological analyses. JS performed laboratory discrimination experiments. BAC performed field discrimination experiments and co-designed the study. JRG co-designed the study, collected specimens, performed microsatellite genotyping, oversaw analysis of EODs and population genetics. All authors contributed to writing the manuscript.

## Acknowledgments

Permits to collect fishes in Gabon and export them for this study were granted by l’Institut de Recherche en Ecologie Tropicale, l’Institut de Recherches Agronomiques et Forestières and the Centre National de la Recherche Scientifique et Technologique. We are grateful for the valuable assistance and logistical support we received from J. D. Mbega and students working in these institutions. All techniques used are in accordance with protocols approved by Cornell University’s Center for Research Animal Resources and Education (CARE). Additionally, we thank C.D. Hopkins (CDH), M. Arnegard, A. McCune and K. Shaw, J. Fetcho, D. Deitcher, S. Mullen as well as two anonymous reviewers for comments on earlier versions of this manuscript. This work was supported by NIMH TG T32 MH015793, NIH TG 2T32GM007469, and NSF 1455405 to JRG, NIH RO1-DC6206, NSF 0818305 to CDH, and NSF 0818390 to BAC.

