## Supporting Information for "Genetic drift does not sufficiently explain patterns of electric signal variation among populations of the mormyrid electric fish *Paramormyrops kingsleyae*"

**Supplementary Figures**

**Figure S1. Example Playback Waveforms: Hybrid EODs**

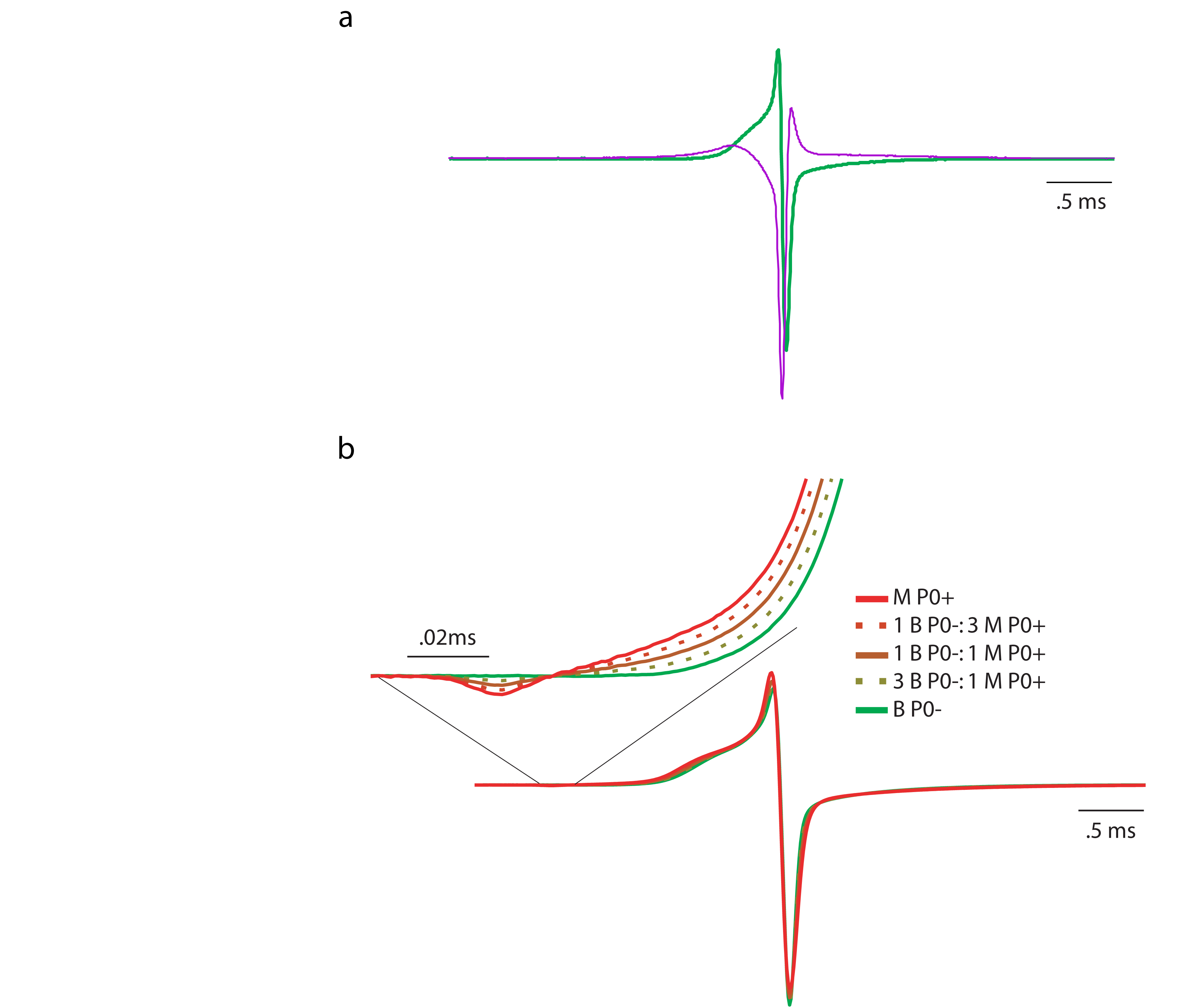
Superimposed, normalized EOD waveforms used in playback experiments, including ‘natural’ EOD waveforms that were P0-present (P0+), P0-absent (P0-) and artificial “hybrid” waveforms (1P0-: 3P0+, 1P0-:1P0+, 3P0-1P0+) made by weighted averages of two natural EODs. Inset expands P0 region x 32.5

**Figure S2. Multidimensional scaling (MDS) plot of EOD waveform variation in 327 *P. kingsleyae* individuals from 11 populations in Gabon.**

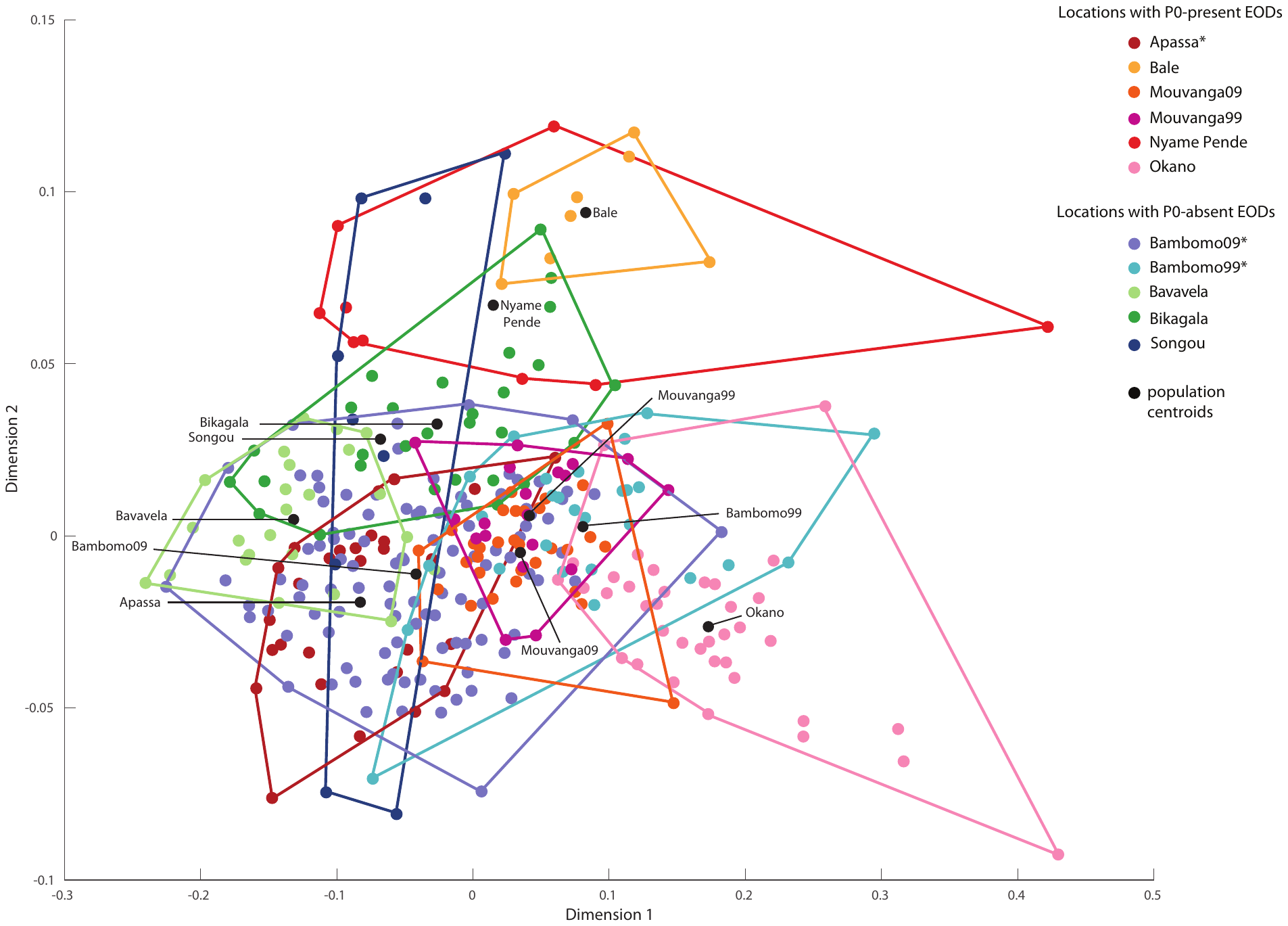

Variation in waveforms was quantified through cross-correlation analysis of signals. Polygons enclose EOD waveforms from each recording locality. Polygon centroids are represented with black dots. Asterisks in the legend represent two populations with mixed signal types (Apassa is mostly composed of P0-present individuals, whereas Bambomo is mostly composed of P0-absent individuals).

**Figure S3. Comparison of inter-population signal distances estimated from PCA (Fig. 3) and from multidimensional scaling of cross-corelated waveforms (Fig. S1).**

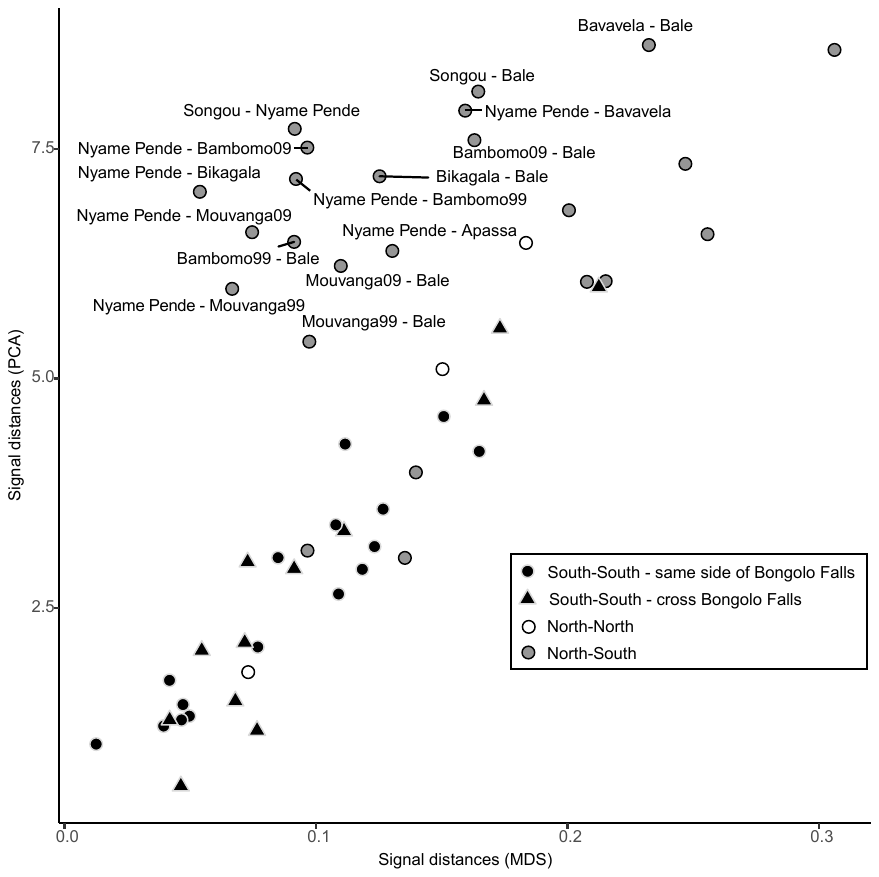

Labels are shown for comparisons where PCA-derived distances exceed MDS-derived ones.

**Figure S4. Mean (and range) of genetic distances for pairwise comparisons of Southern populations separated or on the same side of the Bongolo Falls.**

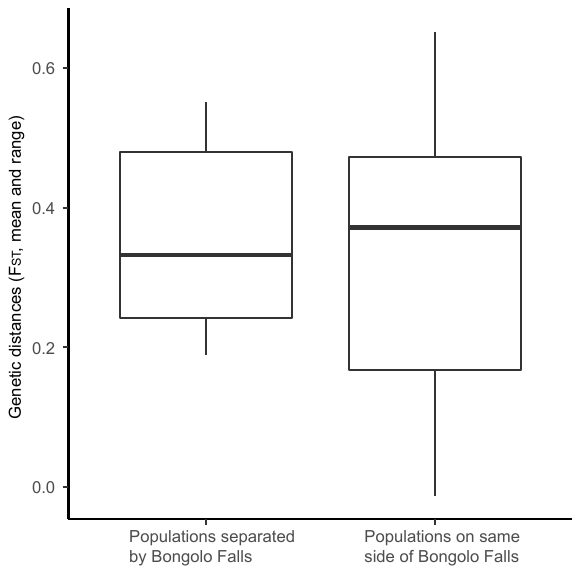

**Figure S5. Correlation plots between genetic, signal, and geographic distances for Southern populations only (n=8 populations).**

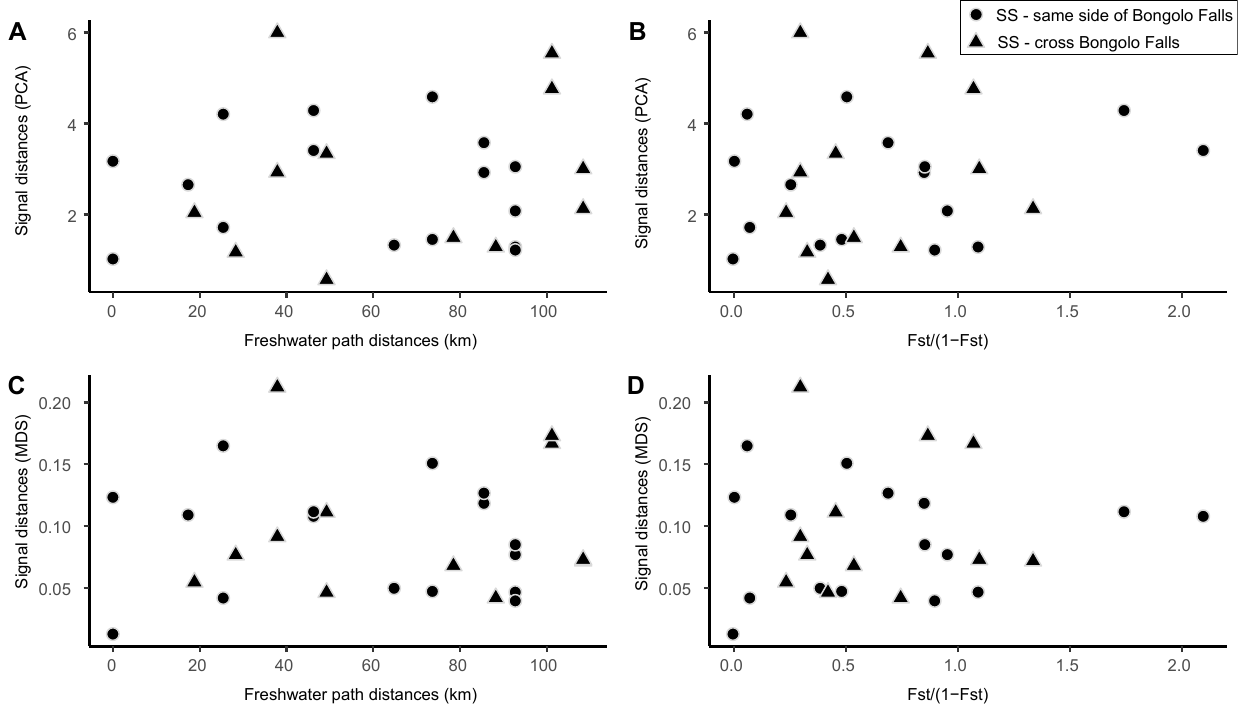

**A.** Correlation between PCA-derived signal and genetic distances. **B.** Correlation between PCA-derived signal and geographic distances. **C**. Correlation between MDS-derived signal and genetic distances. **D.** Correlation between MDS-derived signal and genetic distances. (SS = South-South comparisons).

| **Landmarks** | | |
| --- | --- | --- |
|  | *Name* | *Definition* |
|  | **P0** | Small negative pre-potential; assumed present in all waveforms (re-define this – as the point in the EOD between the beginning of the trace and point P1, with the most negative voltage. |
|  | **P1** | point on the waveform with the Largest head positive voltage |
|  | **P2** | point on the waveform with the Largest head negative voltage |
|  | **S1** | Point on the waveform with the largest first derivative of voltage. slope at the steepest before P1 Point of maximum slope between T1 and P1 (units of measurement?) |
|  | **S2** | Point on waveofmr with minimum first derivative between point P1 and P2 What about point S3 – the point of maximum first derivative after P2 |
|  | **T1** | Waveform beginning; first point to deviate in absolute value from baseline by 2% of peak to peak height |
|  | **T2** | Waveform ending; last point to deviate in absolute value from baseline by more than 2% of peak to peak height |
|  | **ZC1** | Point in waveform closest to the last positive going zero crossing before point P1. voltage crosses baseline from P1-start of record; if exists |
|  | **ZC2** | point in the waveform closest to the first negative going zero crossing between P1 and P2. Point voltage crosses baseline between P1-P2 |
| **Variables** | | |
|  | *Name* | *Definition* |
|  | **aP0** | Area under curve between ZC1-0.5ms and ZC1, (mv • msec) |
|  | **aP1** | Area under curve between ZC2-ZC1 (mv • msec) |
|  | **aP2** | Area under curve between tT2-ZC2 expressed in mv • msec |
|  | **vP0** | Minimum voltage between ZC1-0.5ms and ZC1 (mv) |
|  | **vP1** | Voltage of P1 (mv relative to vP1-vP2=1000 mv) |
|  | **vS1** | Voltage at S1 (mv relative to vP1-vP2=1000 mv ) |
|  | **vS2** | Voltage at S2 (mv relative to vP1-vP2=1000 mv ) |
|  | **tP0** | Time at P0 (msec relative to tP1=0) |
|  | **tP2** | Time at P2 (msec relative to tP1=0) |
|  | **tS1** | Time at S1 (msec relative to tP1=0) |
|  | **tS2** | Time at S2 (msec relative to tP1=0) |
|  | **tZC1** | Time at ZC1 (msec relative to tP1=0) |
|  | **tZC2** | Time at ZC2 (msec relative to tP1=0) |
|  | **Duration** | Total duration (T2-T1, msec) |
|  | **sZC1** | Slope at ZC1 (mv • msec^-1^) |
|  | **sZC2** | Slope at ZC2 (mv • msec^-1^) |
|  | **sS1** | Slope at S1 (mv • msec^-1^) |
|  | **sS2** | Slope at S2 (mv • msec^-1^) |
|  | **fftmax** | Peak frequency of FFT transform of EOD (Hz) |
|  | **fftlo** | Frequency below fftmax at - 3dB (Hz) |
|  | **ffthi** | Frequency above fftmax at - 3dB (Hz) |

**Supplementary Tables**

**Table S2. Landmark and variable definitions.** All variables used in the PCA are defined using these landmarks.

**Table S3. Top ten factor loading values for principal components 1 and 2.**

| **Variable** | **PC1** | **Variable** | **PC2** |
| --- | --- | --- | --- |
| ap2^a^ | -0.325 | vP0^b^ | -0.399 |
| ap1^a^ | 0.320 | fftmax | 0.384 |
| ffthi^a^ | -0.314 | fftlo | 0.366 |
| tZC2 ^a^ | 0.303 | sS2 | 0.288 |
| tP2^a^ | 0.301 | sZC2 | 0.286 |
| tS2^a^ | 0.293 | ap0^b^ | -0.280 |
| duration^a^ | 0.267 | duration | -0.266 |
| vS1 | 0.261 | tP0^b^ | 0.251 |
| sZC2 | 0.208 | tZC1 | 0.247 |
| sS2 | 0.205 | sS1 | -0.227 |

Summary of factor loadings for principal components analysis (Fig. 4). Variables measured in this analysis were described in detail in Table S1.

^a^ Factors for PC1 which are related to overall duration of EOD waveform

^b^ Factors for PC2 which are related to overall magnitude of phase P0.

**Table S4. Results of standard and partial Mantel tests between genetic distances (F_st_), signal distances (obtained through PCA (left) and MDS (right)), geographic distances (km), and presence/absence of Bongolo Falls (Southern populations) at two spatial scales.**

|  |  | PCA-derived signal distances | | |  | MDS-derived signal distances | | |
| --- | --- | --- | --- | --- | --- | --- | --- | --- |
|  |  | *Mantel's r* |  | *p* |  | *Mantel's r* |  | *p* |
| Total range | Correlation |  |  |  |  |  |  |  |
| (n=9 populations) | genetic - geographic | 0.122 |  | 0.209 |  | 0.122 |  | 0.209 |
|  | signal - genetic | 0.067 |  | 0.330 |  | 0.016 |  | 0.455 |
|  | signal - geographic | **0.813** |  | **0.003** |  | **0.369** |  | **0.037** |
|  | Partial correlation |  |  |  |  |  |  |  |
|  | genetic - geographic (regional clusters as covariate) | 0.188 |  | 0.128 |  | 0.188 |  | 0.128 |
|  | signal - genetic (regional clusters as covariate) | 0.020 |  | 0.422 |  | -0.013 |  | 0.503 |
|  | signal - geographic (regional clusters as covariate) | **0.421** |  | **0.030** |  | -0.063 |  | 0.603 |
| Southern populations | Correlation |  |  |  |  |  |  |  |
| (n=6 populations)* | genetic - geographic | **0.775** |  | **0.001** |  | **0.775** |  | **0.001** |
|  | genetic - Bongolo Falls | 0.081 |  | 0.314 |  | 0.081 |  | 0.314 |
|  | signal - genetic | 0.150 |  | 0.201 |  | 0.059 |  | 0.339 |
|  | signal - geographic | 0.069 |  | 0.324 |  | 0.018 |  | 0.365 |
|  | signal - Bongolo Falls | 0.081 |  | 0.314 |  | 0.110 |  | 0.272 |

Significance at the 0.05 level or below is marked in bold.

**Table S5. Results of standard and partial Mantel tests between genetic distances (F_st_/(1-F_st_)), signal distances (obtained through PCA (left) and MDS (right)), geographic distances (km), and presence/absence of Bongolo Falls (Southern populations) at two spatial scales.** Samples from Bambomo and Mouvanga collected from different years are treated as one group for each locality.

|  |  | PCA-derived signal distances | | |  | MDS-derived signal distances | | |
| --- | --- | --- | --- | --- | --- | --- | --- | --- |
|  |  | *Mantel's r* |  | *p* |  | *Mantel's r* |  | *p* |
| Total range | Correlation |  |  |  |  |  |  |  |
| (n=9 populations) | genetic - geographic | -0.041 |  | 0.532 |  | -0.041 |  | 0.532 |
|  | signal - genetic | -0.0004 |  | 0.458 |  | -0.012 |  | 0.468 |
|  | signal - geographic | **0.898** |  | **0.002** |  | **0.480** |  | **0.026** |
|  | Partial correlation |  |  |  |  |  |  |  |
|  | genetic - geographic (regional clusters as covariate) | 0.037 |  | 0.339 |  | 0.037 |  | 0.339 |
|  | signal - genetic (regional clusters as covariate) | 0.087 |  | 0.398 |  | 0.020 |  | 0.398 |
|  | signal - geographic (regional clusters as covariate) | **0.579** |  | **0.024** |  | -0.030 |  | 0.525 |
| Southern populations | Correlation |  |  |  |  |  |  |  |
| (n=6 populations)* | genetic - geographic | **0.536** |  | **0.050** |  | **0.536** |  | **0.050** |
|  | genetic - Bongolo Falls | -0.091 |  | 0.667 |  | -0.091 |  | 0.667 |
|  | signal - genetic | 0.492 |  | 0.111 |  | 0.369 |  | 0.147 |
|  | signal - geographic | 0.164 |  | 0.332 |  | 0.118 |  | 0.365 |
|  | signal - Bongolo Falls | -0.074 |  | 0.600 |  | -0.086 |  | 0.533 |

Significance at the 0.05 level or below is marked in bold. * Significance of these Mantel tests was estimated with the maximum number of permutation possible, (i.e. 719 permutations instead of 1000) due to the reduced size of these distance matrices.

**Table S6. Results of standard and partial Mantel tests between genetic distances (F_st_), signal distances (obtained through PCA (left) and MDS (right)), geographic distances (km), and presence/absence of Bongolo Falls (Southern populations) at two spatial scales.** Samples from Bambomo and Mouvanga collected from different years are treated as one group for each locality.

|  |  | PCA-derived signal distances | | |  | MDS-derived signal distances | | |
| --- | --- | --- | --- | --- | --- | --- | --- | --- |
|  |  | *Mantel's r* |  | *p* |  | *Mantel's r* |  | *p* |
| Total range | Correlation |  |  |  |  |  |  |  |
| (n=9 populations) | genetic - geographic | 0.114 |  | 0.283 |  | 0.114 |  | 0.283 |
|  | signal - genetic | 0.106 |  | 0.321 |  | 0.056 |  | 0.371 |
|  | signal - geographic | **0.898** |  | **0.002** |  | **0.480** |  | **0.026** |
|  | Partial correlation |  |  |  |  |  |  |  |
|  | genetic - geographic (regional clusters as covariate) | 0.114 |  | 0.292 |  | 0.114 |  | 0.292 |
|  | signal - genetic (regional clusters as covariate) | 0.071 |  | 0.378 |  | 0.017 |  | 0.402 |
|  | signal - geographic (regional clusters as covariate) | **0.579** |  | **0.024** |  | -0.030 |  | 0.525 |
| Southern populations | Correlation |  |  |  |  |  |  |  |
| (n=6 populations)* | genetic - geographic | **0.719** |  | **0.001** |  | **0.719** |  | **0.001** |
|  | genetic - Bongolo Falls | -0.005 |  | 0.533 |  | -0.005 |  | 0.533 |
|  | signal - genetic | 0.424 |  | 0.161 |  | 0.337 |  | 0.176 |
|  | signal - geographic | 0.164 |  | 0.332 |  | 0.118 |  | 0.365 |
|  | signal - Bongolo Falls | -0.074 |  | 0.600 |  | -0.086 |  | 0.533 |

Significance at the 0.05 level or below is marked in bold. *Significance of these Mantel tests was estimated with the maximum number of permutation possible, (i.e. 719 permutations instead of 1000) due to the reduced size of these distance matrices.
