## Appendices for "Genetic drift does not sufficiently explain patterns of electric signal variation among populations of the mormyrid electric fish *Paramormyrops kingsleyae*"

**Appendix 1: Temperature correction on signal variation quantified through multidimensional scaling of signal cross correlations**

Kramer and Westby (1985) showed that EOD durations are stretched and compressed due to temperature-dependent processes with a Q_10_ factor of 1.49 in the mormyrid *Gnathonemous petersii*. In our collections, recording temperature varied from 20.5 to 25.1°C. Therefore, we standardized all recorded waveforms to the mean temperature of our dataset (23.5°C) using a Q_10_ of 1.49 in the following way. Regrettably, temperature data was not available for three waveforms (two from Bambomo09, one from Bikagala), which were thus excluded from this analysis (n=324).

1. We adjusted the EOD time base of each waveform by multiplying the A/D digitizer’s sampling rate by factor *f*, using the following equation:

$$f={10}^{\frac{23.5^{\circ}C-Tobs}{10}}log(Q10)$$

where Q10=1.49 and T_obs_ corresponds to the temperature in the recording chamber at the time of the EOD recording.

2. As cross-correlation analysis involves the progressive sliding of one waveform past another, we could not simply correct each waveform by changing its time base to compress or stretch it, but instead needed to directly adjust the voltage values of each waveform to reflect our Q_10_ correction. To do so, we interpolated the voltage values of each EOD at the pre-adjusted time base points from the tracings of the EOD at its corrected time base using the *interp1* function in MATLAB.
3. Cross-correlation of all temperature corrected EODs was then performed as described in the main text, and multidimensional scaling was applied to the cross-correlation matrix using the *mdscale* function in MATLAB with Kruskal’s normalized stress 1 criterion (Kruskal and Wish 1978). The number of dimensions was set to N=2 which resulted in a stress of 0.0505, considered to give a good ordination representation with low probability of misinterpretation (Clarke 1993). Figure A1 represents the multidimensional scaling plot of Q10 corrected waveforms.

The shift of points in multidimensional scaling space between temperature uncorrected (Fig. S2) and corrected signals (Fig. A1) is essentially focused on Songou individuals, with very little change to the configuration of all other population polygons. In order to estimate whether temperature correction affected our evaluation of the potentials of different shaping electric signal variation, we performed the same set of Mantel tests described in the main text on the signal distances estimated from the MDS plot of Q_10_ corrected signals. We calculated the electric signal distance between population centroids as the Euclidean distance between group centroids in the Q_10_ corrected MDS space using the R package *vegan.* We found that Mantel tests on signal distances estimated from temperature corrected signals were entirely consistent with Mantel test results from temperature-uncorrected signals: the only slightly significant correlation was found to be between signal and geographic distance throughout the whole range of Gabon (Mantel rho=0.25, p=0.05, Table A1). We conclude from these analyses that temperature did not have a significant effect on *P. kingsleyae* signal variation in our study. While temperature correction did have a detectable effect in terms of the MDS configuration, this effect was negligible compared to the variation between individuals and between localities sampled. We therefore report analyses and results in the main text without any temperature correction.

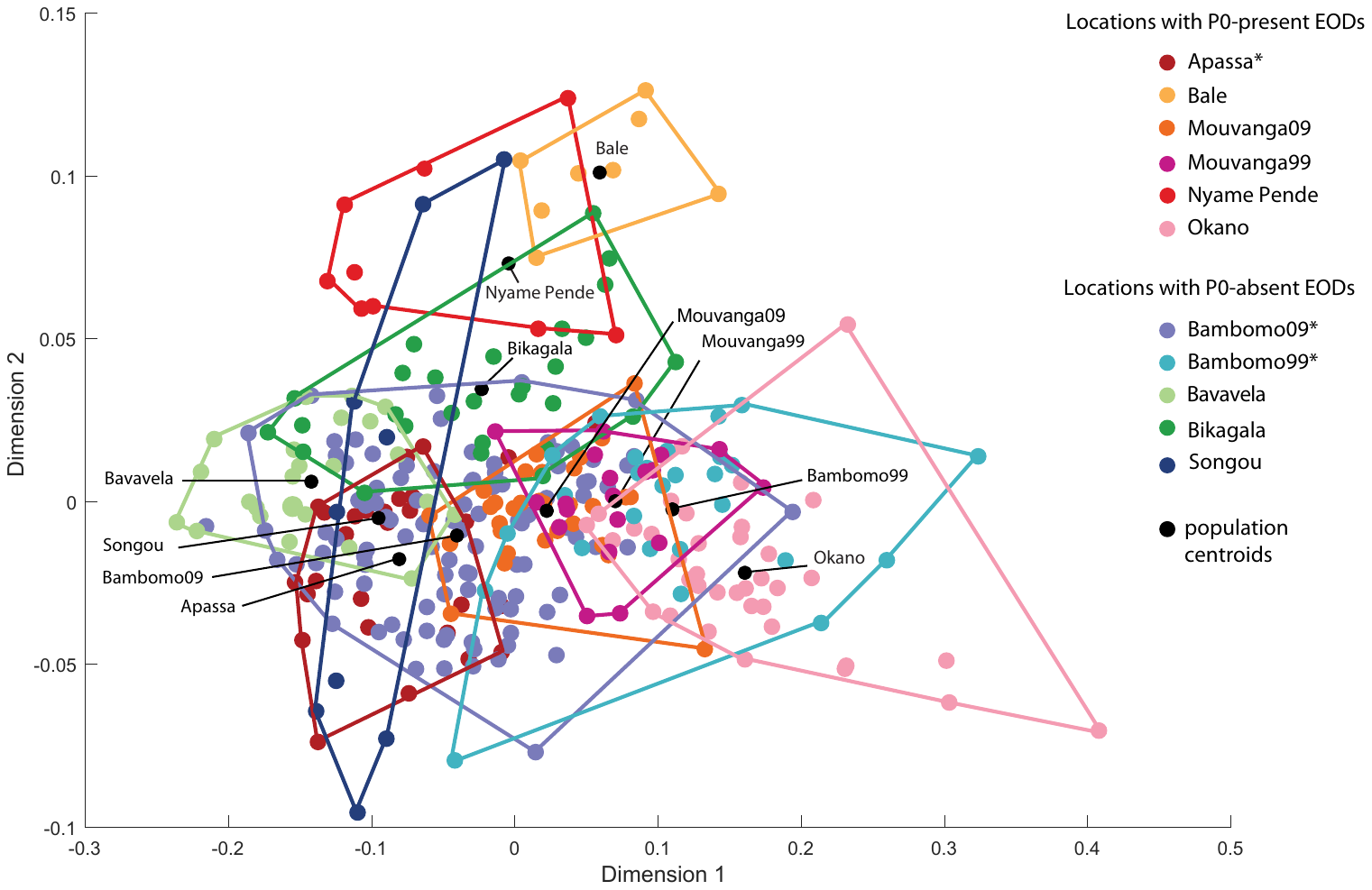

**Figure A1.** Multidimensional scaling (MDS) plot of Q_10_ corrected EOD waveform variation in 324 *P. kingsleyae* individuals from 11 populations in Gabon. Variation in waveforms was quantified through cross-correlation analysis of signals (see Carlson et al. (2011)). Polygons enclose EOD waveforms from each recording locality. Polygon centroids are represented with black edges. Asterisks in the legend represent two populations with mixed signal types (Apassa is mostly composed of P0-present individuals, whereas Bambomo is mostly composed of P0-absent individuals).

**Table A1. Results of standard and partial Mantel tests between genetic distances (F_st_/(1-F_st_)), temperature corrected signal distances (obtained through MDS), geographic distances (km), and presence/absence of Bongolo Falls (Southern populations) at two spatial scales.**

|  |  | MDS-derived signal distances | | |
| --- | --- | --- | --- | --- |
| Group |  | *Mantel's r* |  | *p* |
| Total range | Correlation |  |  |  |
| (n=11 populations) | signal - genetic | -0.013 |  | 0.522 |
|  | signal - geographic | **0.255** |  | **0.042** |
|  | Partial correlation |  |  |  |
|  | signal - genetic (regional clusters as covariate) | 0.007 |  | 0.432 |
|  | signal - geographic (regional clusters as covariate) | -0.020 |  | 0.518 |
| Southern populations | Correlation |  |  |  |
| (n=8 populations) | signal - genetic | 0.062 |  | 0.330 |
|  | signal - geographic | -0.040 |  | 0.529 |
|  | signal - Bongolo Falls | 0.067 |  | 0.331 |

Significance at the 0.05 level or below is marked in bold.

### **Appendix 2: Power analysis of microsatellites for inferring genetic differentiation**

We performed power analysis simulations using powsim version 4.1 (Ryman and Palm 2006) to determine if our sample sizes, number of microsatellite loci, and allele diversity were sufficient to detect genetic differentiation. This analysis simulates sampling from *s* populations of equal effective population size N_e_ that have split from a base population through random drift for *t* generations to an expected predefined level of differentiation (measured as F_ST_ = 1 – (1 – 1/2N_e_)^t^). Samples are then drawn from the simulated populations and used for testing genetic homogeneity using Fisher’s exact test based on all loci simultaneously. We used empirical sample sizes to simulate 10000 random sets of 11 populations with expected F_ST_ values of 0.001 – 0.05. Since effective population sizes (N_e_) for this system are unknown, we used a combination of effective population size (Ne=500; Ne=1000; Ne=5000) and time since divergence estimates. Power estimates were obtained as the proportion of tests that indicated significant differentiation.

These analyses showed that our sample sizes and specific genetic markers were adequate for detecting levels of genetic differentiation as low as F_ST_ = 0.003 with a high probability (>0.9) (Fig. A2). We can therefore consider our genetic markers as sufficiently powerful for the purpose of this study, namely to give a reliable estimate of genetic differentiation between *P. kingsleyae* allopatric populations. Indeed, the lowest significant F_st_ value detected in our dataset was 0.055, well above the F_ST_ power threshold of 0.003 calculated in POWSIM. Among all pairwise differentiation tests, three were considered non-significant, namely Mouvanga Creek (1999 vs. 2009 F_ST_=-0.013), Bambomo Creek (1999 vs. 2009 F_ST_=0.001), and Bale Creek vs. Nyamé Pendé Creek, F_ST_=0.01). Concerning the Mouvanga and Bambomo comparisons, it is possible that this was due to a lack of power (F_st_ estimates are lower than the calculated F_ST_ power threshold of 0.003).

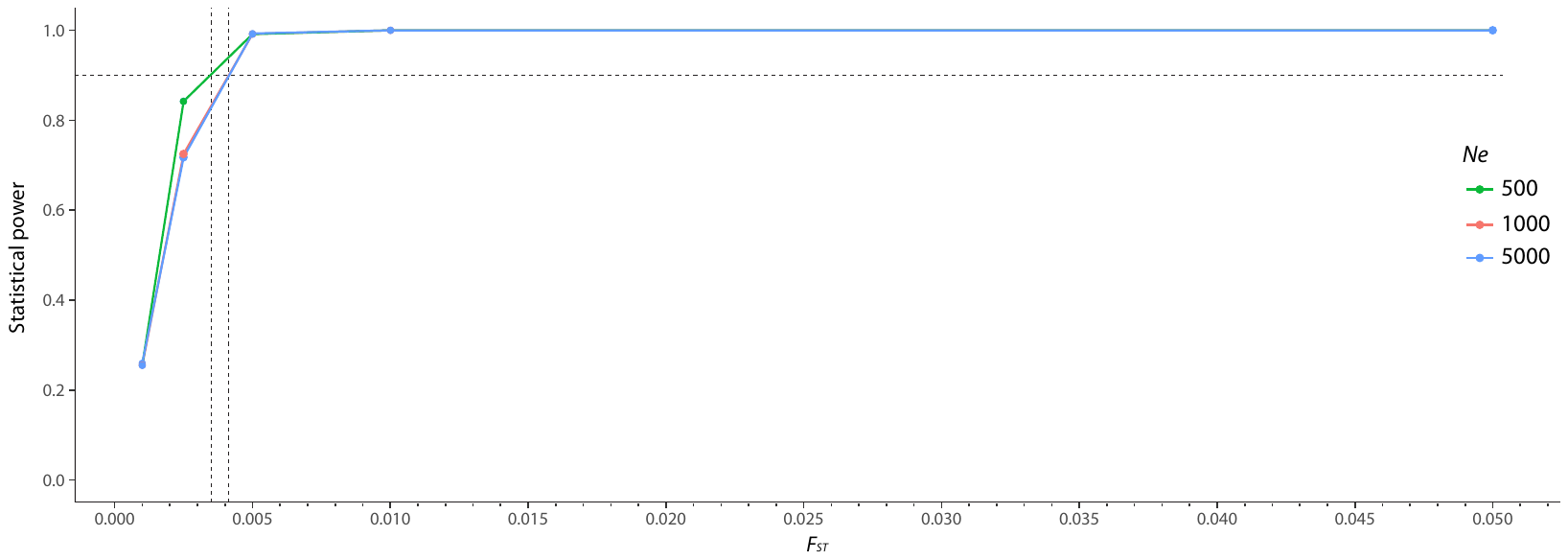

**Figure A2** Power analysis simulations for obtaining significant outcomes in tests of genetic differentiation, using five microsatellite loci and the specific marker characteristics and sample sizes of this study. Simulations were performed using powsim (Ryman and Palm 2006). The dotted line indicates the minimum level of genetic differentiation that can be detected with 90% statistical power. *N*e indicates the effective population size used in different simulation runs.
